# Single nucleus multi-omics regulatory atlas of the murine pituitary

**DOI:** 10.1101/2020.06.06.138024

**Authors:** Frederique Ruf-Zamojski, Zidong Zhang, Michel Zamojski, Gregory R. Smith, Natalia Mendelev, Hanqing Liu, German Nudelman, Mika Moriwaki, Hanna Pincas, Rosa Gomez Castanon, Venugopalan D. Nair, Nitish Seenarine, Mary Anne S. Amper, Xiang Zhou, Luisina Ongaro, Chirine Toufaily, Gauthier Schang, Joseph R. Nery, Anna Bartlett, Andrew Aldridge, Nimisha Jain, Gwen V. Childs, Olga G. Troyanskaya, Joseph R. Ecker, Judith L. Turgeon, Corrine K. Welt, Daniel J. Bernard, Stuart C. Sealfon

**Author notes:** Co-corresponding authors **Corresponding authors:** Frederique Ruf-Zamojski, Stuart C. Sealfon.

## Abstract

The pituitary regulates growth, reproduction and other endocrine systems. To investigate transcriptional network epigenetic mechanisms, we generated paired single nucleus (sn) transcriptome and chromatin accessibility profiles in single mouse pituitaries and genome-wide sn methylation datasets. Our analysis provided insight into cell type epigenetics, regulatory circuit and gene control mechanisms. Latent variable pathway analysis detected corresponding transcriptome and chromatin accessibility programs showing both inter-sexual and inter-individual variation. Multi-omics analysis of gene regulatory networks identified cell type-specific regulons whose composition and function were shaped by the promoter accessibility state of target genes. Co-accessibility analysis comprehensively identified putative cis-regulatory regions, including a domain 17kb upstream of *Fshb* that overlapped the fertility-linked rs11031006 human polymorphism. *In vitro* CRISPR-deletion at this locus increased *Fshb* levels, supporting this domain’s inferred regulatory role. The sn pituitary multi-omics atlas (snpituitaryatlas.princeton.edu) is a public resource for elucidating cell type-specific gene regulatory mechanisms and principles of transcription circuit control.

## INTRODUCTION

The pituitary gland plays a critical role in the regulation of the key physiological functions, as it integrates regulation of the central nervous system with that of the endocrine system. It consists of an anterior, an intermediate, and a posterior lobe. The anterior lobe, which comprises ~80% of the gland, contains five major hormone-producing cell types (somatotropes, gonadotropes, lactotropes, thyrotropes, and corticotropes), as well as non-endocrine cells. Recent single cell (sc) transcriptome studies highlighted a heterogeneity in pituitary cell populations and sub-populations, and other molecular signatures that may reflect different functional cell states^1, 2, 3, 4, 5^. Sc epigenetic assays, particularly methylome analysis, are more definitely distinguishing distinct cell types from transient cell states^6^, and provide insight into epigenetic regulatory mechanisms.

Gene regulatory programs are generally orchestrated by transcription factors (TFs) via interaction with cis-regulatory genomic DNA sequences located in or around target genes. Epigenetic mechanisms, including changes in chromatin accessibility and DNA methylation (for review^7^), play crucial roles in the regulation of gene expression. Epigenomic profiling technologies have been developed to explore various layers of epigenetic regulation at sc resolution, including single nucleus (sn) ATACseq, which measures chromatin accessibility, and genome-wide mapping of DNA methylation^8, 9^. The integration of sc epigenomics with sc transcriptomics provides the opportunity to elucidate the regulatory programs and epigenetic mechanisms underlying cell type-specific gene expression, and to resolve intercellular heterogeneity^10^.

One limitation of sc sequencing technologies is that tissue dissociation can elicit artifactual gene expression^11, 12, 13, 14^. Unlike sc-based methods, sn approaches are compatible with snap-frozen tissue samples and minimize *ex vivo* expression changes^15^. The transcriptome complexity identified by snRNAseq is comparable to that of scRNAseq methods^16^. Besides its reliability for profiling the transcriptome at sc resolution^11, 13, 17, 18, 19^, sn isolation also allows mapping of the chromatin-regulatory landscape^20, 21, 22^ and genome-wide measurement of DNA methylation^23^.

In the present work, we sought to resolve transcriptional regulatory mechanisms in murine pituitary cells and identify cellular heterogeneity at transcriptomic and epigenomic levels. We analyzed over 65,000 nuclei from individual snap-frozen male and female pituitaries for a parallel analysis of their transcriptome and genome-wide chromatin accessibility profiles. For DNA methylation profiling, we pooled 30 snap-frozen male pituitaries. We showed how the resulting sn multi-omics atlas provides insight into regulatory network and gene control mechanisms that are relevant to physiology and disease.

## RESULTS

### Sn multi-omic profiling in murine pituitaries

We determined at sn resolution transcriptome and chromatin accessibility landscape in snap-frozen adult male and female murine pituitaries, and DNA methylation status in pooled male murine snap-frozen pituitaries (**Fig. 1)**. On the day of the assay, nuclei were isolated from individual pituitaries, and joint snRNAseq and snATACseq assays performed on each individual animal sample. From six animals, 35,707 nuclei were assayed by snRNAseq assays and 33,443 by snATACseq (**Supplementary Figure 1, Supplementary Table 1**). 5-methylcytosine sequencing 2 assay (snmC-seq2^23^) was performed on 2,756 nuclei from a pool of 30 male pituitaries (**Supplementary Figure 1, Supplementary Table 1**).

**Figure 1:**
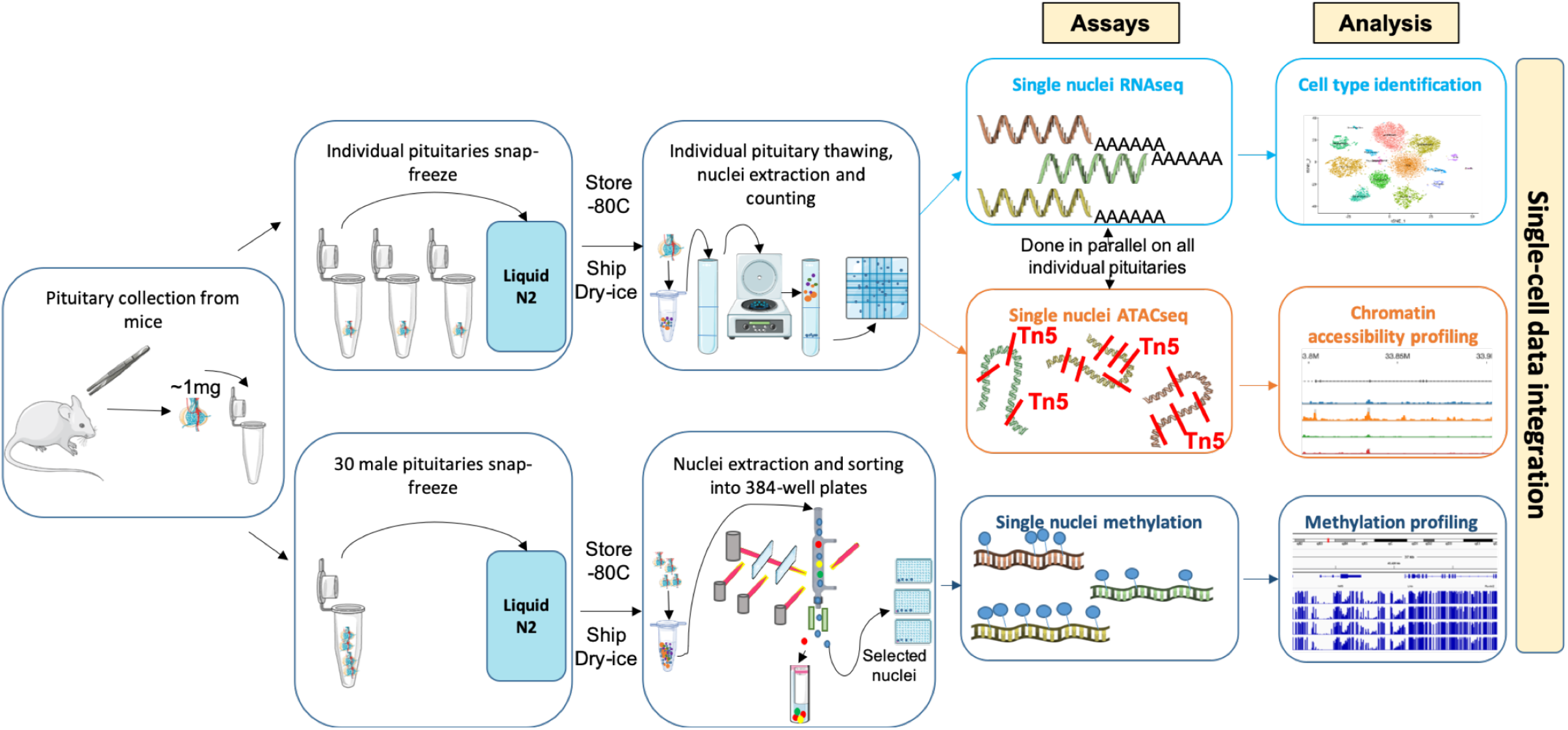
Overview of pituitary sn multi-omics experimental design. SnRNAseq and snATACseq assays were performed simultaneously on each individual snap frozen murine pituitary. For sn methylation assays, pituitaries were pooled from 30 male mice. Additional studies were performed on cryopreserved dissociated murine pituitary cells, including scRNAseq, snRNAseq, and snATACseq (see **Supplementary Figures 1-2**)

Additional datasets were generated using alternate tissue handling, processing and assay protocols (**Supplementary Figure 1, Supplementary Table 1**, see **Supplementary Methods**). These included different pituitary dissociation protocols followed by pituitary cell cryopreservation, as well as sc vs. snRNA sequencing methods. Overall, snap-frozen pituitaries and gently dissociated pituitaries elicited the highest quality sequencing data regardless of the assay type (**Supplementary Figure 1**). Our analysis focused predominantly on data from snap-frozen pituitaries, given their high quality and the minimization of *ex vivo* changes.

### SnRNAseq analysis identifies the major pituitary cell types

An average of 6,000 nuclei per snap-frozen pituitary were analyzed by snRNAseq, detecting about 2,000 genes per nucleus. Cells were clustered using Seurat’s shared nearest nuclei approach, annotated per cluster and visualized using t-Stochastic Neighbor Embedding representation (tSNE). The ranking of differentially expressed genes in each cluster relative to all other cells (**Supplementary Figures 2-3**), combined with the distribution of key pituitary marker allowed annotation of the major pituitary cell type clusters in males (**Fig. 2a,b**) and females (**Supplementary Figure 2**). Besides the major pituitary cell types, we also identified stem (progenitor) cells, proliferating cells, pituicytes, and cell types found in and around blood vessels (macrophages, endothelial cells, pericytes; **Fig. 2a, Supplementary Figure 2**). Interestingly, Seurat clustering analysis of male pituitaries distinguished three contiguous regions within the somatotrope cluster (**Fig. 2a** and **Supplementary Figure 4**). Several transcripts distinguished the two poles of the overall somatotrope cluster, including *Lhfpl3*, *Egfeml*, *Dpp10* and *B3galt1* for Somatotrope 1 (Som1), and *Tcerg1l*, *Bach2* and *Dgkg* for Somatotrope 2 (Som2; see **Supplementary Figure 4** for a complete list of genes). The third somatotrope region, Somatotrope 0 (Som0), appeared closer to the lactotropes but was only seen in males. The absence of Som0 cells in females might be the result of fewer nuclei being sequenced by snRNAseq in the female samples. Most Som0 cells expressed *Gh* and *Sgcz*, with varying but not significant levels of *Prl*, and represented about 5% of all cells in our dataset (**Fig. 2a,b**). Som 0 was unlikely to include doublets because the absolute number of transcripts (UMI counts) was on par with that of the other somatotropes (**Fig. 2c**). Mitochondrial gene content in the Som0 region was low, excluding dying cells (**Fig. 2d**). The gene expression pattern in the Som0 region was distinct from Som 1/2, lactotropes, and a cluster of lactotrope/somatotrope doublets (**Supplementary Figure 4**).

**Figure 2:**
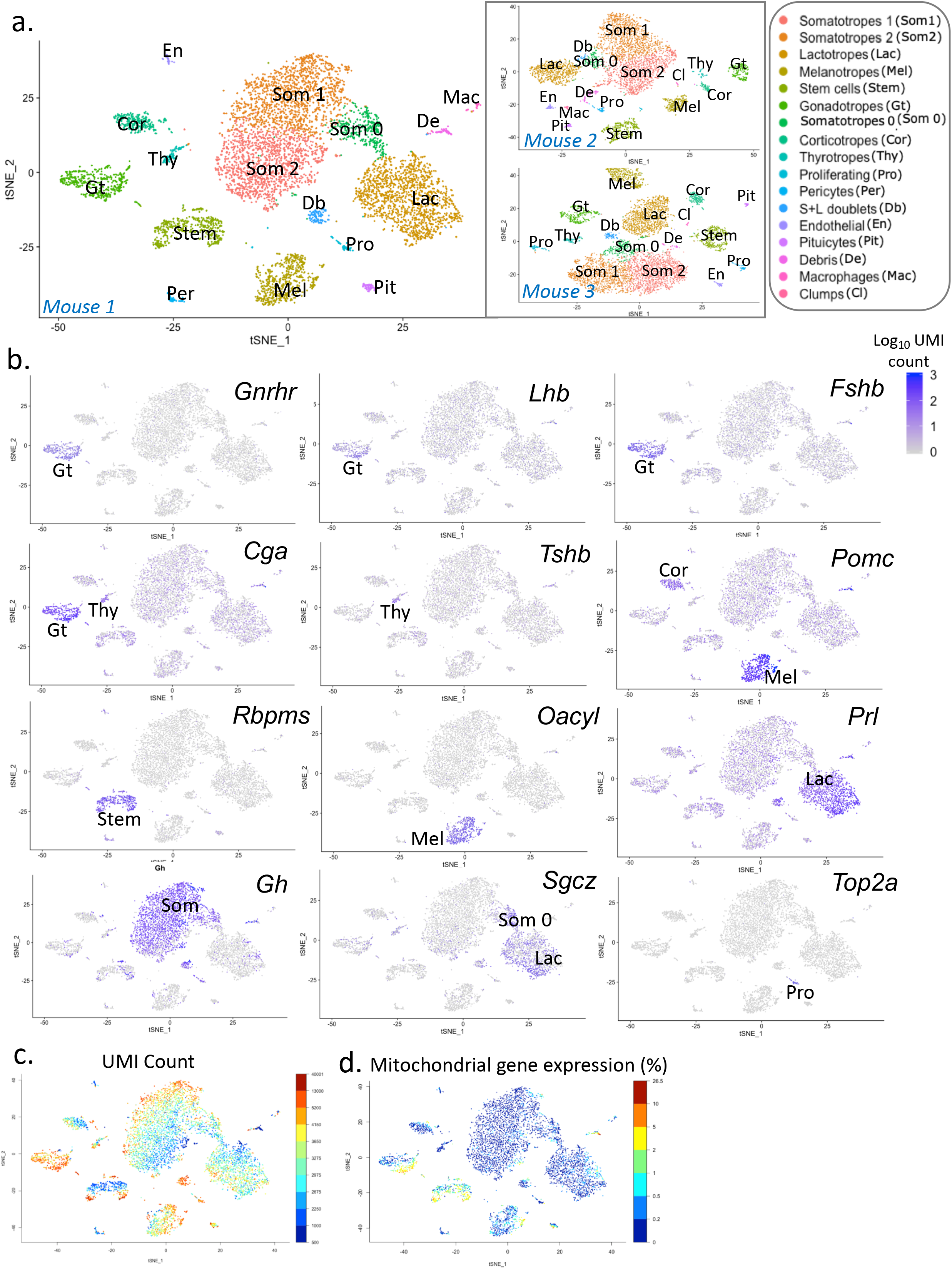
Identification of mouse pituitary cell types by sn transcriptomic analysis. **a**. tSNE representation of sn transcript expression in individual snap-frozen individual male pituitaries with cells colored by type. Shown are data derived from 3 individual male pituitaries. **b**. Individual feature plots for transcripts used to identify major cell types in one male pituitary. **c**. UMI count graph showing the number of individual transcripts in one male pituitary. **d**. Graph showing mitochondrial gene expression in one male pituitary. **b-d** show the same male pituitary (Mouse 1).

Analysis of scRNAseq from dissociated cells also identified the major classical hormone-producing cell types (**Supplementary Figure 5**). Clusters with low numbers of transcripts most likely corresponded to non-viable cells after cell dissociation (**Supplementary Figure 5**) as they were undetected in snRNAseq from either dissociated (**Supplementary Figure 6**) or snap-frozen pituitaries (**Fig. 2**, **Supplementary Figures 2-3**).

### Confirmation of cell type identity by sn epigenome analysis

We employed snATACseq to analyze chromatin accessibility in the same male and female pituitaries analyzed by snRNAseq. Data generated from 65,411 analyzed nuclei from snap-frozen pituitaries (**Fig. 3, Supplementary Figure 8**) or cryopreserved dissociated pituitaries (**Supplementary Figures 7, 11b**) exhibited a canonical fragment-size distribution with clearly resolved mono- and multi-nucleosomal modes, as well as a high signal-to-noise ratio at transcription start sites (TSS; **Supplementary Figure 8b**). TSS enrichment scores ranged from 9.08 to 11.82, with an average of 10 for all 6 snap-frozen pituitary samples (**Supplementary Figure 8b**). An average of 5,000 nuclei were analyzed per pituitary sample, with an average of 30,000 peaks per nucleus (**Supplementary Figure 8**). Unbiased genome-wide analysis of chromatin accessibility resulted in the identification of 10-11 cell type clusters in male samples (**Fig. 3A**). In female samples, we identified 13-15 cell type clusters (**Supplementary Figure 8a**), likely because more nuclei were sequenced in these samples by snATACseq, enabling the detection of additional clusters. Cluster identity was assigned (**Fig. 3a**) based on measurements of open chromatin (i.e. peaks of accumulated reads) at the promoters of major cell type marker genes (**Fig. 4**). The results validated the major pituitary cell types determined by snRNAseq analysis (**Fig. 2a**), including the gradient of somatotropes (**Fig. 3a**, **Supplementary Figure 9**). The genes whose promoter accessibility differentiated the two poles of the somatotrope cluster included *Ngb* and *Rem1* for Som1, and *Ascl2* and *Oit1* for Som2 (**Supplementary Figure 9**).

**Figure 3:**
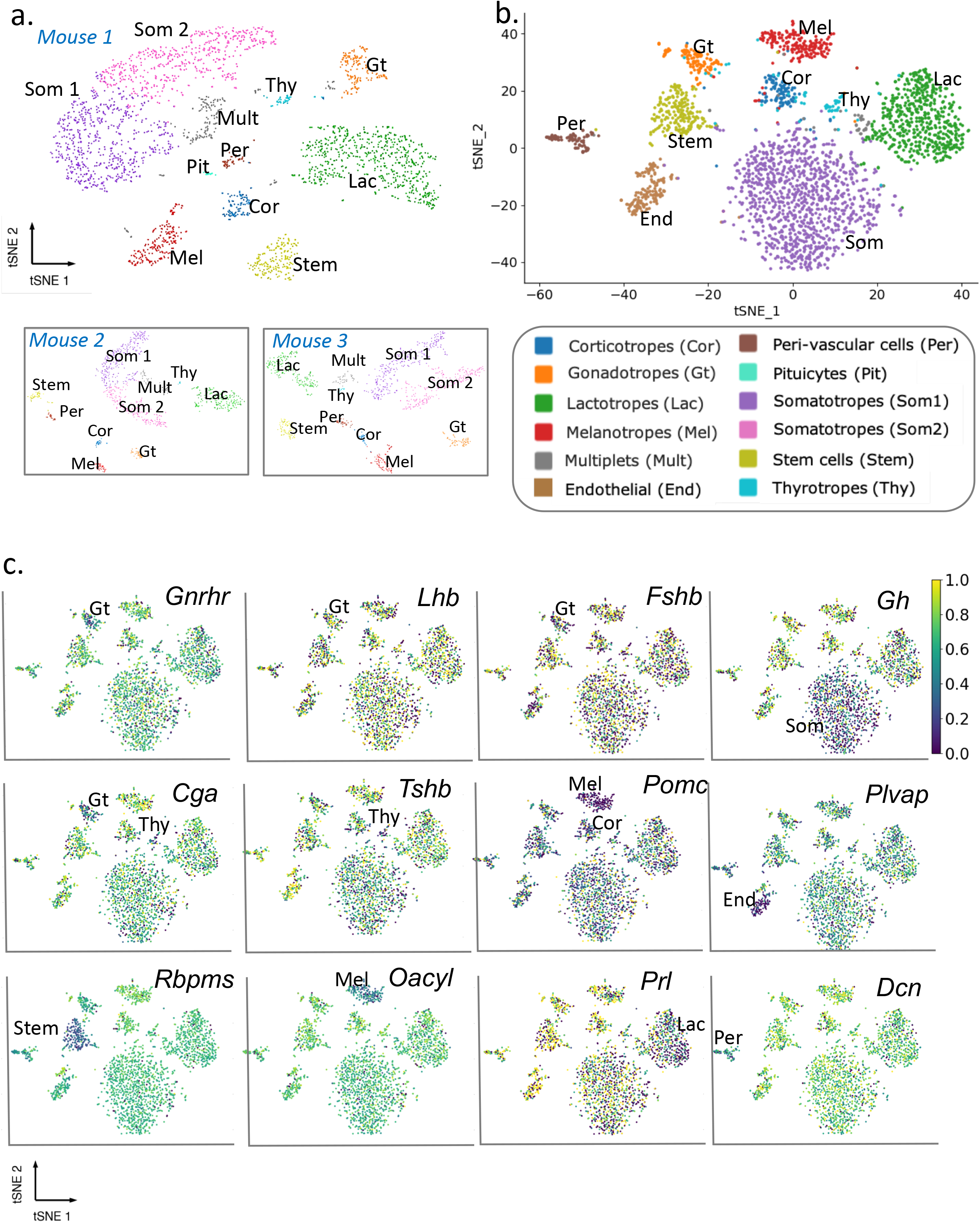
Identification of mouse pituitary cell types by sn epigenomic analysis. **a**. tSNE representation of sn chromatin accessibility in individual snap-frozen male pituitaries with cells colored by types. Shown are data from the same 3 individual male pituitaries presented in **Fig. 2**. **b**. tSNE representation of snDNA methylation from a pool of 30 snap-frozen male pituitaries with cells colored by type. **c**. Feature plot showing DNA methylation status of the genes indicated. Hypomethylation is associated with higher transcript expression (compare with **Fig. 2b**).

**Figure 4:**
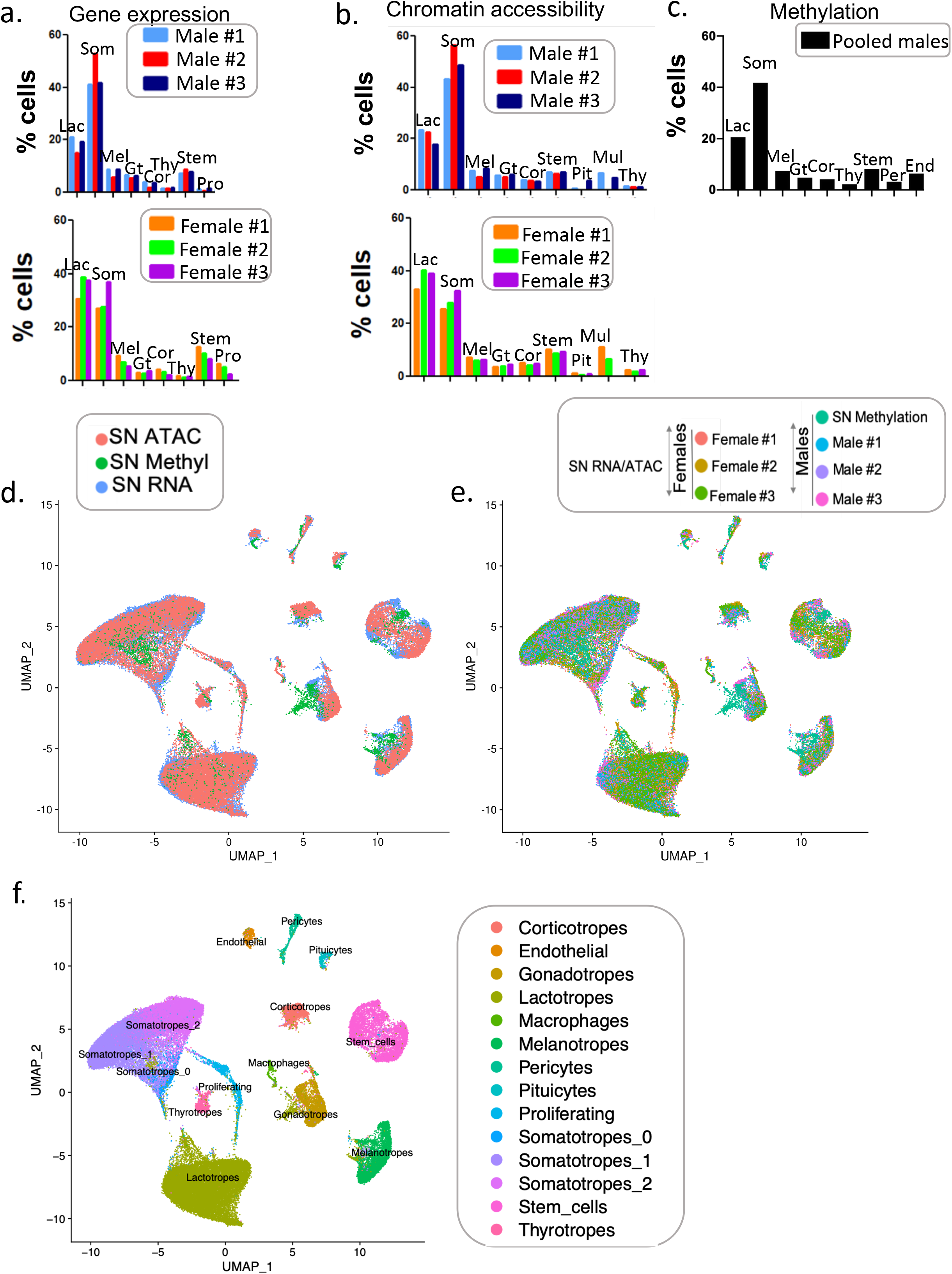
Multi-omics pituitary cell type integration. **a,b**. Cell type proportions identified in each animal from snRNAseq (**a**) and snATACseq (**b**) datasets show individual differences that are concordant (See also **Supplementary Table 2**). Differences between sexes were significant for lactotropes and somatotropes (p<0.0001) for both assays. **c**. Cell type proportions identified in sn methylation data from pooled male pituitaries. **d**. UMAP overlay of the snATACseq (red), sn methylation (green), and snRNAseq (blue) datasets. **e**. UMAP of the overlay of all samples with the three omics modalities. The snRNAseq and snATACseq data are color-coded by animal. The male sn methylation data are also included. **f**. UMAP showing the cluster identification per cell type based on the snRNAseq annotations.

To correlate open chromatin sites with DNA methylation profiles and gain further insight into the epigenomic state of pituitary cells, we performed sn methylation assay on 2,756 nuclei isolated from 30 pooled male pituitaries. Genome-wide analysis of CG DNA methylation levels (mCG) enabled the detection of 9 cell clusters (**Fig. 3b**). Cell cluster identity was defined based on the depletion of mCG levels at the promoters of known pituitary cell type markers. Non-CG DNA methylation (mCH) was a rare event in the pituitary sample, and thus was not used for cell type assignment. The identity of most endocrine cell type clusters was clearly established, as well as that of stem/progenitor cells, endothelial cells, and pericytes, as their methylation patterns were restricted to single groups of cells (**Fig. 3b,c**). As expected, there was an inverse relationship between gene expression level (**Fig. 2b**) and DNA methylation level of the corresponding promoter (**Fig. 3c**), suggesting that promoters with low methylation levels were more likely to be accessible and transcribed. Cell type identification was more challenging for the gonadotrope and thyrotrope clusters (the latter containing only 49 cells), requiring the use of additional gene markers (*Cga*, *Tshb*, *Gnrhr* besides *Fshb* and *Lhb* for gonadotropes; *Tshb* and *Cga* for thyrotropes). In contrast with snRNAseq and snATACseq, sn methylation data analysis did not identify a gradient of somatotropes, suggesting that these transcriptome changes may represent cell states rather than cell types. Overall, the major pituitary cell types identified correlated well with those found by both snRNAseq (**Fig. 2a**) and snATACseq (**Fig. 3a**) analysis.

### Murine pituitary multi-omics sn atlas

Determination of cell type proportions in the snRNAseq data (**Fig. 4a, Supplementary Table 2a**) revealed that females had significantly more lactotropes and fewer somatotropes than males (**Supplementary Table 2**). Similar cell type proportions were inferred from the snATACseq data obtained from the same individual pituitaries (**Fig. 4b**, **Supplementary Table 2b**) and corresponded to those obtained by sn methylation in a sample of pooled male pituitaries (**Fig. 4c**, **Supplementary Table 2c**). Generally, male-female differences in cell type proportions identified by either gene expression or promoter accessibility were in agreement, providing further support for the correct identification of cell types across different assay modalities.

Using the Seurat data integration pipeline, we overlaid snRNAseq, snATACseq, and sn methylation data (**Fig. 4d-f**), following the assumption that higher chromatin accessibility and lower methylation levels would result in higher gene expression. Genome-wide chromatin accessibility and methylation data were converted to gene-level data and integrated to the transcriptomic data by using a nearest neighbors analysis. All modalities fit well with each other for pituitary cell type identification (**Fig. 4d,f**). Additionally, data from individual animals also corresponded well (**Fig. 4e,f**). Markers from the snRNAseq analysis distinguished 17 different cell clusters (**Fig. 4f**, **Fig. 2a**), while fewer clusters were detected by snATACseq and sn methylation (11 and 9 clusters, respectively; **Figs. 3**, **Supplementary Figures 2**,**8**).

We confirmed the identity and assignment of the main pituitary cell types by integrating the data obtained by snATAC, sn methylation, and snRNAseq for eight major pituitary markers (**Fig. 5**). For instance, in gonadotrope cells, the promoters of *Cga* (**Fig. 5a**), *Lhb* (**Fig. 5h**), and *Fshb* (**Fig. 5i**) are accessible, hypomethylated, and these cells express *Cga* (**Fig. 5a**), *Lhb* (**Fig. 5h**), and *Fshb* (**Fig. 5i**) mRNAs. Although no differentially methylated regions (DMR) were identified in the Lhb promoter, we detected a promoter region (red box, **Fig. 5h**) whose mCG level was lower in gonadotropes relative to other cell types. Interestingly, we discovered an open chromatin region upstream of the Fshb promoter (red box, **Fig. 5i**). The other pituitary cell types were similarly confirmed based on chromatin accessibility, methylation profile, and transcript levels of pituitary markers (**Fig. 5**).

**Figure 5:**
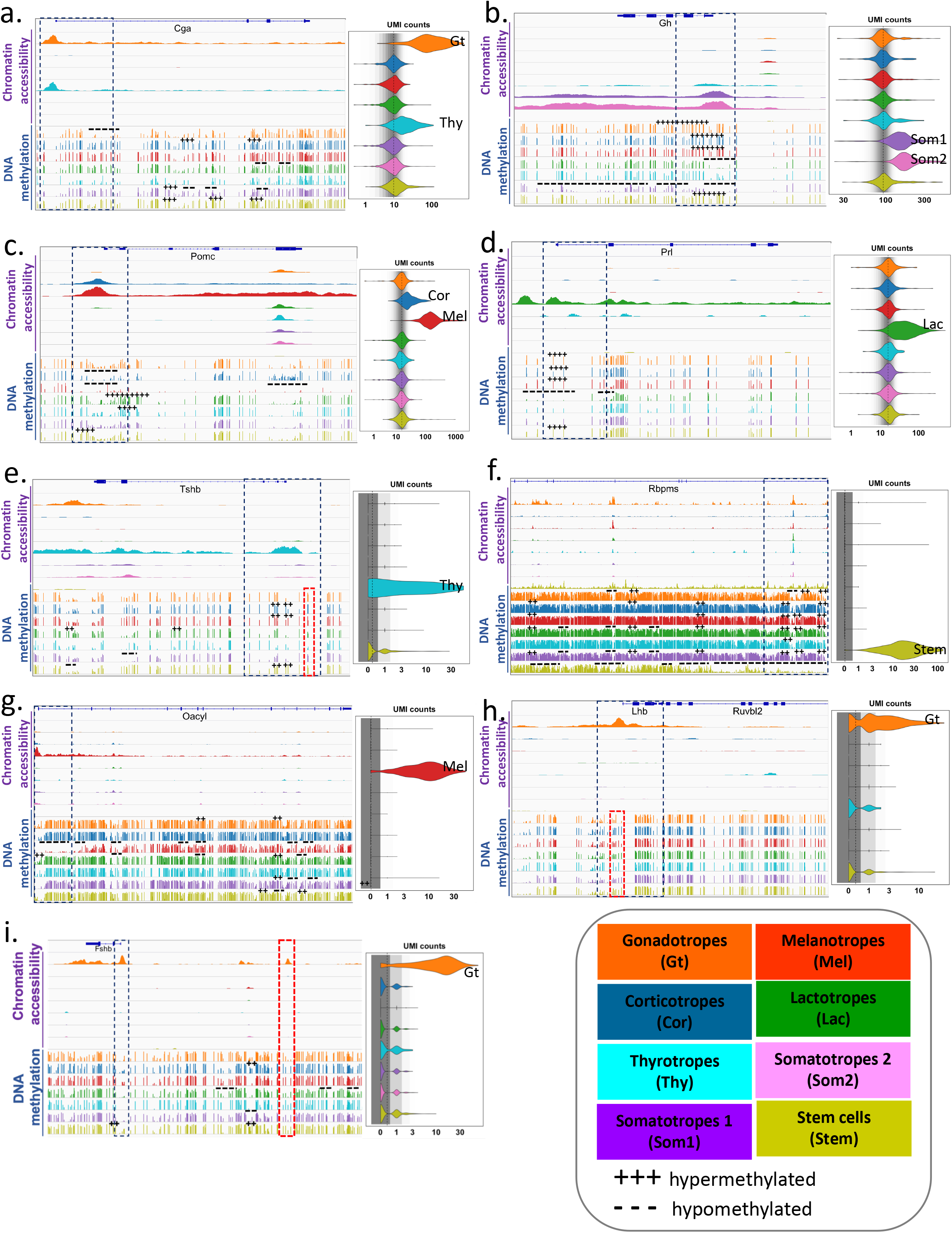
Multi-omics state of representative pituitary genes by cell type. **a-i.** Shown by cell type are genome browser tracks for chromatin accessibility (top), DNA methylation (bottom), and violin plots of transcript expression for the indicated genes. The dotted boxes denote the area around the promoter region of each gene. Red dotted boxes highlight either differentially methylated regions (**e,h**) and/or open chromatin region (**i**) for the indicated genes. For the DNA methylation, the differentially methylated regions (DMR) are shown as hypermethylated (+++), or hypomethylated (−−−).

To enable researchers to navigate our sn mouse pituitary multi-omics atlas, we created a web-based portal accessible at snpituitaryatlas.princeton.edu.

### Transcriptional and epigenetic program inter-animal variation

To characterize coordinated gene expression and accessibility programs, we used the Pathway Level Information ExtractoR framework (PLIER^24^). When applied to RNA expression or chromatin accessibility at gene promoters, PLIER identified sets of genes (Latent Variable, LV) that vary in concert across different cell types or samples. PLIER uses known pathways and other gene set annotations to improve identification of these LV sets and to associate them with biological processes. LVs are analogous to eigengenes^25^ and can provide a summary level of overall expression or accessibility per cell type or sample, which can be displayed as heatmaps or bar graphs (See **Fig. 6**, **Supplementary Figures 10-11**).

**Figure 6:**
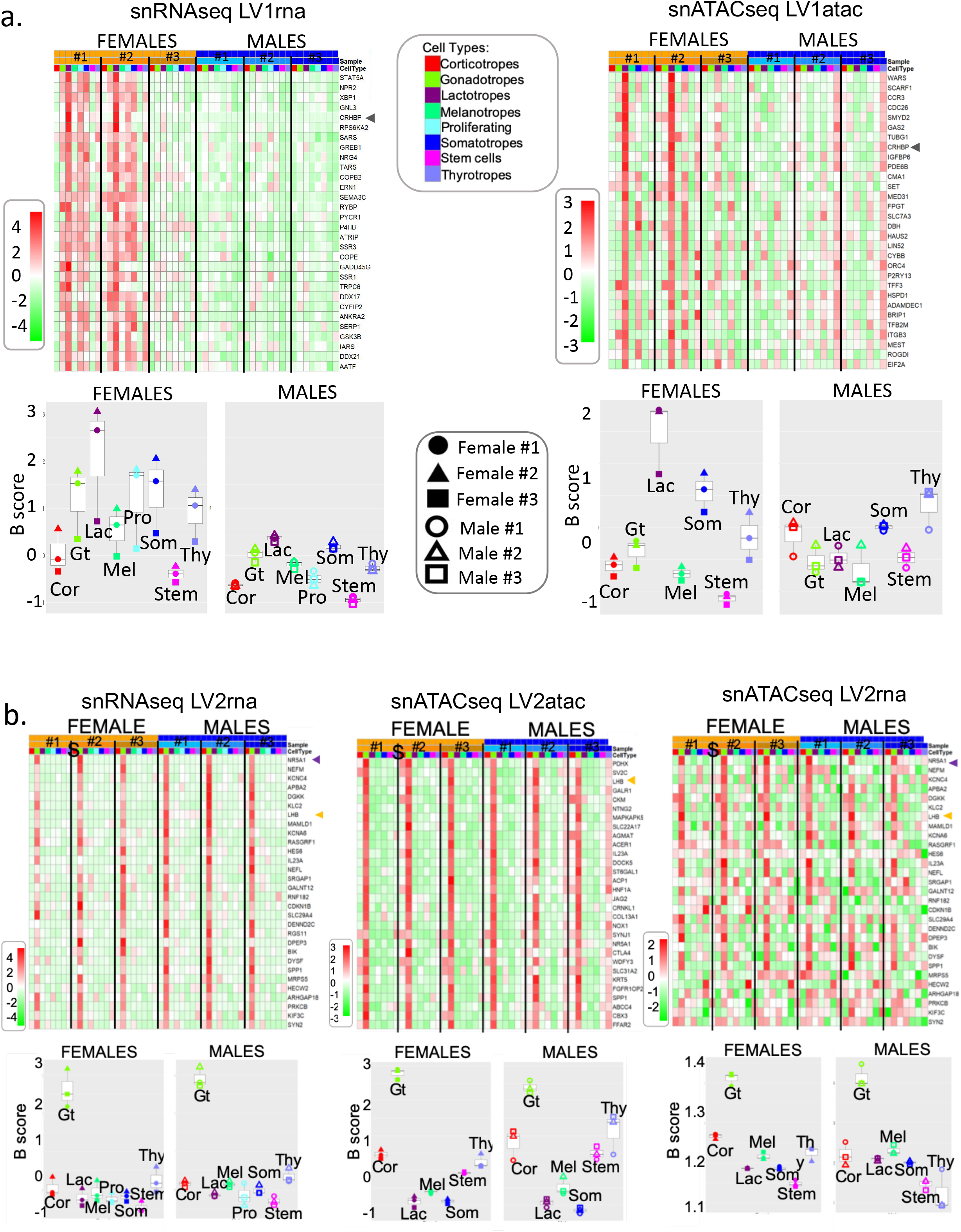
Inter-animal differences in gene expression and chromatin accessibility. **a**. Identification of latent variables (LVs) showing sex and inter-animal differences using PLIER. LV1rna (left panel) and LV1atac (right panel) were inferred from the snRNAseq and snATACseq data, respectively. Depicted are a heatmap (top) and expression levels (bottom) of each LV across samples and cell types. Differences between sexes were significant for LV1rna (p= 1.1e^−5^, Wilcoxon, all cell types), and for LV1atac (p= 0.02 for lactotropes only). A top driver gene for LV1rna and LV1tacat is indicated with an arrow. **b**. Heatmap and expression levels of an LV that is elevated in gonadotropes. The LV was identified from the snRNAseq data (left panel, LV2rna), applied to snATAC data (right panel, LV2rna), and a similar LV was identified from the snATAC data (center panel, LV2atac). Selected genes that are characteristic of gonadotropes are indicated with arrows.

PLIER analysis of the snRNAseq and snATACseq datasets identified LVs showing differential expression between sexes or predominant expression in each of the major pituitary cell types (**Supplementary Figure 10**). For example, LV1rna and LV1atac (**Fig. 6a**) showed increased expression in females (Wilcoxon p= 1.1e^−5^) and in female lactotropes (Wilcoxon p=0.02) respectively. Comparing LV1rna in an independent scRNAseq dataset from dissociated male and female pituitaries confirmed the differential expression between females and males (**Supplementary Figure 11**, Wilcoxon p=1.1e^−8^). In snap-frozen pituitaries, LV1rna and LV1atac also showed similar patterns of expression in individual animals, with one female pituitary exhibiting a male-like pattern for each LV (**Fig. 6a**). This female also showed apparent differences in cell type ratios determined by snRNAseq, with 37% somatotropes vs. 27.3% and 27.7% in the other two females (**Supplementary Table 2**). Correct sex identification of all animals was confirmed by *Xist* and Y chromosome gene expression, as well as open chromatin accessibility in the Y chromosome for the males. We speculated that the female showing a male-like pattern of expression might be a non-cycling female. Particularly, one of the top drivers for LV1rna and LV1atac was *Crhbp*, an estrogen-regulated gene (arrows**, Fig. 6a**). The expression and accessibility levels of the *Crhbp* gene, and the expression of the estrogen-responsive *Greb1* gene (involved in the estrogen receptor-regulated pathway^26, 27, 28, 29^) were much higher in the other two females than in the female showing a male-like pattern (**Fig. 6a**). *Crhbp* gene expression was previously reported to be more elevated in females relative to males and to increase during proestrus^30^. While these animals were randomly selected with no determination of cycle status, the LV analysis suggests the identification of a stage where estradiol is low in this female, consistent with an absence of cycling.

Many of the genes in LV1rna and promoter accessibility sites in LV1atac do not correspond. This is expected as these two LVs were identified by independent PLIER deconvolution of the two data types. Nonetheless, the formulation that LV1rna and LV1atac captured related transcriptional and epigenetic pathways is supported by their similar patterns of variation between sexes and among individual animals, as well as their significant overlap of transcripts and proximal gene accessibility sites (hypergeometric test for overlap of 17 from the first 200 features; p= 3.6e^−5^).

Predominant coordinated gene and chromatin accessibility programs were identified for all major pituitary cell types (**Supplementary Figure 10**). Gonadotrope-predominant LVs are shown in **Fig. 6b**. Transcriptome-based LV2rna was elevated in gonadotropes (p=1.4e^−5^, Kruskal-Wallis test) and showed a similar expression level in all animals (**Fig. 6b**, left panel). Gene accessibility-based LV2atac showed a similar pattern of increased gonadotrope expression (p=1.5e^−6^) among all animals (**Fig. 6b**, middle panel). The overlap between LV2rna and LV2atac genes was highly significant (hypergeometric test for overlap of 31 from the first 200 features; p=6.1e^−15^), supporting the formulation that they may capture biologically related transcriptional and epigenetic regulatory programs. Finally, when applying the expression pattern of LV2rna directly to the snATACseq data, we also found a pattern of increased gonadotrope expression (**Fig. 6b**, right panel; p=2.8e^−5^). Overall, we identified inter-sexual and inter-individual differences in coordinated gene expression and promoter accessibility programs, as well as coordinated cell type-specific programs.

### Epigenetic control of cell type-specific regulons

We next sought to characterize the individual TF-driven units (regulons^31^) of the pituitary gene regulatory networks (GRN) using SCENIC^32^. This computational approach identified 344 pituitary regulons varying between 3 to 4,761 genes with a median size of 50 genes per regulon. Determination of regulon activity in each cell type showed clusters of regulons with higher activity in different cell types (**Fig. 7a, Table 1,** snpituitaryatlas.princeton.edu). Many regulons were associated with driver TFs characteristic of pituitary cell types. For example, regulons showing high activity in the gonadotropes included TFs that are involved in gonadotrope development and/or regulation of gonadotropin subunit gene expression (*Gata2*, *Foxl2*, *Nr5a1*, *Pitx1*, *Smad3*), or are gonadotrope-enriched (*Foxp2*; for review^1, 33, 34, 35^). We examined the ability of this differential cell type regulon expression to classify cells by projecting the regulon activities within each cell onto Uniform Manifold Approximation and Projection (UMAP^36^) axes. Notably, this projection of regulon activities recapitulated the major cell type clusters identified by Seurat analysis of sn transcript expression (**Fig. 7b,** compare with **Fig. 2a**). Regulon-based clustering separated lactotropes into three clusters. Annotation of these clusters by individual animal showed that the clustering was based on differences in lactotrope regulon activity between the two sexes and among individual female pituitaries (**Fig. 7c**). The identification of regulon differences across sex and individual animals is consonant with the pattern of variation found in PLIER analysis-derived LVs (**Fig. 6a**).

**Figure 7:**
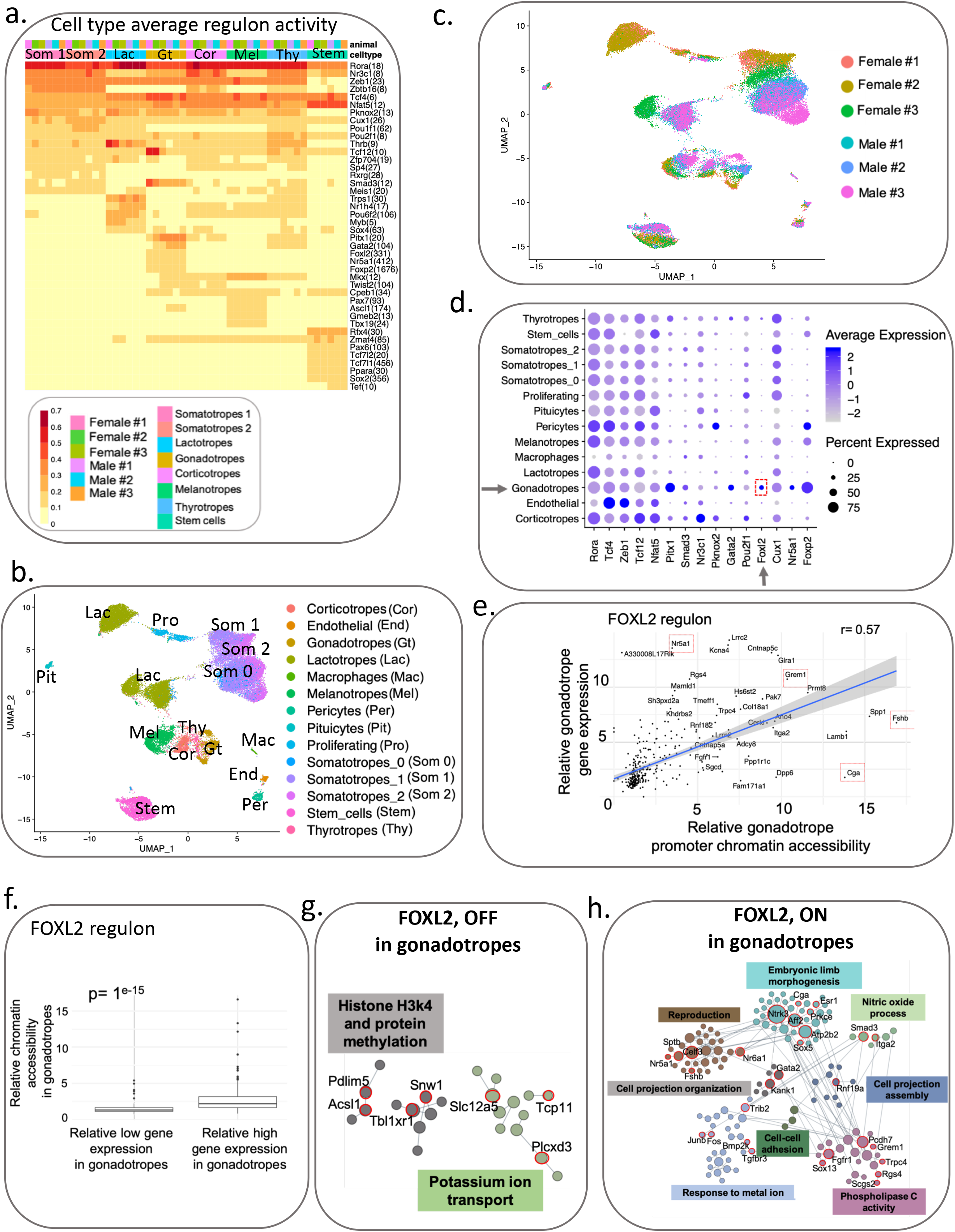
Gonadotrope-specific regulon activity. **a**. Heatmap of the SCENIC cell type average regulon activity scores per animal. **b**. UMAP and cell type identification from the regulon analysis. **c**. UMAP of the individual animal samples overlaid on the cell clusters obtained with the SCENIC analysis. **d**. Top regulons in male gonadotropes and their relative expression in other cell types. **e**. Scatter plot and correlation of the 331 genes composing the FOXL2 regulon based on their relative gene expression and relative TSS chromatin accessibility in male gonadotropes. Selected genes are boxed in red. **f**. Boxplots of the relative chromatin accessibility within gene bodies for the FOXL2 regulon separated into relatively low expression vs. relatively high expression in male gonadotropes. **g**. Annotated gene modules based on FOXL2 regulon genes that are not expressed (off) in male gonadotropes. **h**. Annotated gene modules based on FOXL2 regulon genes expressed (on) in male gonadotropes. Panels **d-h** are based on data from 3 individual male samples. See **Supplementary Figure 13** for corresponding analysis of female samples and **Supplementary Figure 14** for another gonadotrope-specific regulon NR5A1.

**Table 1:** SCENIC highest activity regulons per cell type.

| Cell type |  |  |  |  |  |  |  |  |  |  |  |  |
| --- | --- | --- | --- | --- | --- | --- | --- | --- | --- | --- | --- | --- |
| **Somatotrope 1** | Rora | Nr3c1 | Zeb 1 | Zbtb16 | Tcf4 | Nfat5 | Pknox2 | Cux1 | Pou1f1 | Pou2f1 | Thrb | Tcf12 |
| **Somatotrope 2** | Rora | Nr3c1 | Zeb 1 | Zbtb16 | Nfat5 | Tcf4 | Cux1 | Pou1f1 | Thrb | Tcf12 | Pou2f1 | Smad3 |
| **Lactotropes** | Rora | Zeb1 | Thrb | Tcf4 | Trps1 | Pknox2 | Nfat5 | Nr1h4 | Pou6f2 | Myb | Pou1f1 | Pou2f1 |
| **Gonadotropes** | Rora | Tcf4 | Zeb 1 | Tcf12 | Nfat5 | Pitx1 | Smad3 | Nr3c1 | Pknox2 | Gata2 | Pou2f1 | Foxl2 |
| **Corticotropes** | Rora | Nr3c1 | Zeb1 | Tcf4 | Nfat5 | Pknox2 | Zbtb16 | Pou2f1 | Tcf12 | Mkx | Cux1 | Twist2 |
| **Melanotropes** | Rora | Tcf4 | Zeb 1 | Nfat5 | Mkx | Pax7 | Pou2f1 | Tcf12 | Ascl1 | Cux1 | Gmeb2 | Tbx19 |
| **Thyrotropes** | Rora | Zeb1 | Tcf4 | Nr3c1 | Nfat5 | Thrb | Pknox2 | Pou2f1 | Tcf12 | Cux1 | Pitx1 | Pou1f1 |
| **Stem cells** | Nfat5 | Tcf4 | Rora | Rfx4 | Nr3c1 | Zmat4 | Pax6 | Pou2f1 | Cpeb1 | Tcf12 | Tcf7l2 | Tcf7l2 |

Because regulon composition is inferred from gene expression in all cell types, it may overestimate the set of genes activated within a specific cell type. In principle, regulon activity and composition in each cell type are subject to expression of the driver TF within each cell type. We investigated the hypothesis that regulon composition could be restricted by the epigenetic state of its target genes. We initially examined the cell type-specific expression of the TFs driving each regulon, especially in the gonadotropes (**Fig. 7d**). To test whether epigenetic mechanisms may contribute to cell type-specific regulon expression we first studied the FOXL2 regulon. Among the 331 genes identified by SCENIC as members of the FOXL2 regulon, only ~200 were expressed in the gonadotropes. The relative gonadotrope levels of gene expression and promoter chromatin accessibility for all FOXL2 regulon target genes were highly correlated in males (**Fig. 7e**) and in females (**Supplementary Figure 13**). Generally, genes without open promoters were not transcribed (**Fig. 7e**, lower left corner), whereas genes with open promoters were transcribed, with varying levels of gene expression. Gene body accessibility was significantly greater for regulon genes with higher relative expression in the gonadotropes compared to those with lower relative expression (p=1.0e-15, **Fig. 7f**). To determine whether this differential chromatin accessibility correlated with DNA methylation changes, we examined the promoter methylation levels of the regulon genes data. While relative methylation levels of the regulon genes showing high vs. low relative expression in the gonadotropes were not significant (p=0.6), comparison of the highest and lowest expressed genes yielded statistical significance (p=0.008; **Supplementary Figure 12**). These results indicate that promoter accessibility is a fundamental determinant of activity of FOXL2 regulon genes within the gonadotrope, whereas methylation has a more limited influence.

To examine functional consequences of this epigenetic control of the FOXL2 regulon expression in the gonadotropes, we compared the processes represented by the regulon targets that are poorly expressed (off) vs. those that are turned on in the gonadotropes. This functional network module analysis demonstrated that regulon targets that are off in gonadotropes encompass different biological functions (**Fig. 7g**) than the active targets (**Fig. 7h**).

To determine whether promoter accessibility was a general mechanism for cell type-specific regulons, we characterized two other regulons. Like *Foxl2*, *Nr5a1* expression was largely restricted to gonadotropes (**Fig. 7d**). Promoter accessibility was a major determinant of the composition and functional pathways of the NR5A1 regulon in gonadotropes (**Supplementary Figure 14**). The correlation between regulon gene expression and promoter and gene body methylation levels was not significant (data not shown). We next examined the FOXP2 regulon, which was expressed in several pituitary cell types (**Fig. 7d**). The gene composition of this regulon was dramatically different in gonadotropes, melanotropes, and stem cells (**Fig. 8a**). Notably, the correlation between cell type specific levels of gene expression and promoter chromatin accessibility was high in all three cell types (**Fig. 8b,c**; female gonadotropes **Supplementary Figure 15**), yet the FOXP2 regulon functional annotations differed in each cell types (**Fig. 8d**). Altogether, these results support the formulation that the regulon composition and functional processes at cell type resolution are shaped by the epigenetic state of targets in that cell type. We propose a model for the three layers of control (driver expression, target site presence, gene accessibility) that influence the composition of cell type specific regulons and their associated functional processes (**Fig. 8e**).

**Figure 8:**
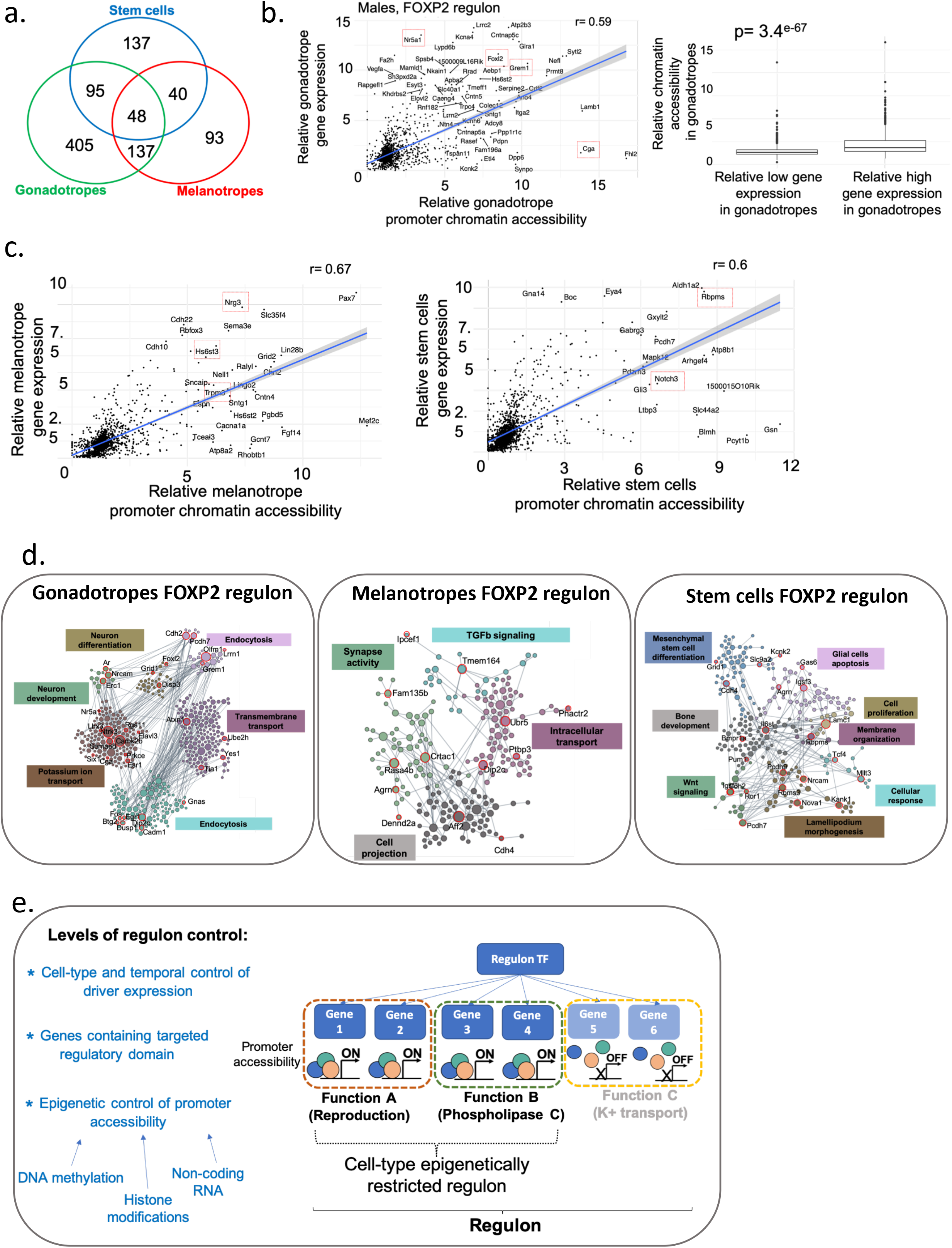
Model of cell type-specific regulon control mechanism. **a**. Venn diagram showing which FOXP2 regulon genes are expressed in other cell types. **b**. Left panel, scatter plot and correlation of the 1676 genes composing the FOXP2 regulon showing relative gene expression and relative TSS chromatin accessibility in male gonadotropes. Right panel, boxplot of the relative chromatin accessibility within gene bodies for the FOXP2 regulon comparing relatively low expressing and relatively high expressing genes in gonadotropes. **c**. Scatter plots and correlations of the FOXP2 regulon based on their relative gene expression and relative TSS chromatin accessibility in male melanotropes (left panel) and stem cells (right panel). Characteristic melanotrope and stem cell genes are boxed in red. **d**. Annotated gene modules showing the differences in function of the FOXP2 regulon in gonadotropes, melanotropes, and stem cells. **e**. Model of the mechanisms proposed for cell type-specific regulation of regulon composition and function. All analyses are based on combined male samples. For corresponding analysis of female samples see **Supplementary Figure 15**.

### CRISPR validation of a co-accessible domain upstream of *Fshb*

Patterns of co-accessibility between distal elements and their target promoters can be utilized to build a genome-wide map of cis-regulatory sequences from sc data^37^. We applied Cicero to our snATACseq data to identify putative regulatory regions of individual genes. Co-accessible regions of classical pituitary gene markers were found within specific cell types (**Supplementary Figure 16,** snpituitaryatlas.princeton.edu). As an example, we focused on co-accessible domains upstream of the *Fshb* promoter, identifying several potential regions of interest (**Fig. 9a, Supplementary Figure 16**). One gonadotrope-specific co-accessible region was located 17 kb upstream of the *Fshb* promoter. The sequence of this putative regulatory region corresponded to the human single nucleotide polymorphism (SNP) rs11031006 (**Fig. 9a**, boxed region; see also **Fig. 5i**) which has been linked to alterations in female fertility^38, 39^. Multispecies comparative sequence analysis (PhyloP(60), PhastCons^40, 41^) showed high sequence conservation of this domain across 60 vertebrate species (**Fig. 9b,c**). Together, co-accessibility analysis, sequence conservation and a fertility-linked human SNP support the hypothesis that this upstream domain contributes to the regulation of *Fshb*. To test this hypothesis, we introduced CRISPR deletions in this region in the murine LβT2 gonadotrope cell line. *Fshb* expression and FSH secretion were undetectable in the parent gonadotrope cell line at baseline and following GnRH/activin A treatment. In contrast, deletion of the TTATTT sequence in this domain (**Fig. 9c**) led to an increase in baseline *Fshb* expression (**Fig. 9d**). Increased *Fshb* expression and FSH secretion were also observed in this mutant following GnRH/activin A treatment. Two additional CRISPR modifications at this site (either a T Deletion/ TATT insertion; or a TTT deletion) also led to measurable baseline *Fshb* expression levels, albeit lower than that observed in the TTATTT deletion line. These findings suggest that the reproductive phenotype associated with human SNP rs11031006 results from altered regulatory control of the *FSHB* gene via this co-accessible upstream domain, and support the value of co-accessibility analysis, available on the atlas web portal, for the identification of putative cis-regulatory domains for any genes of interest.

**Figure 9:**
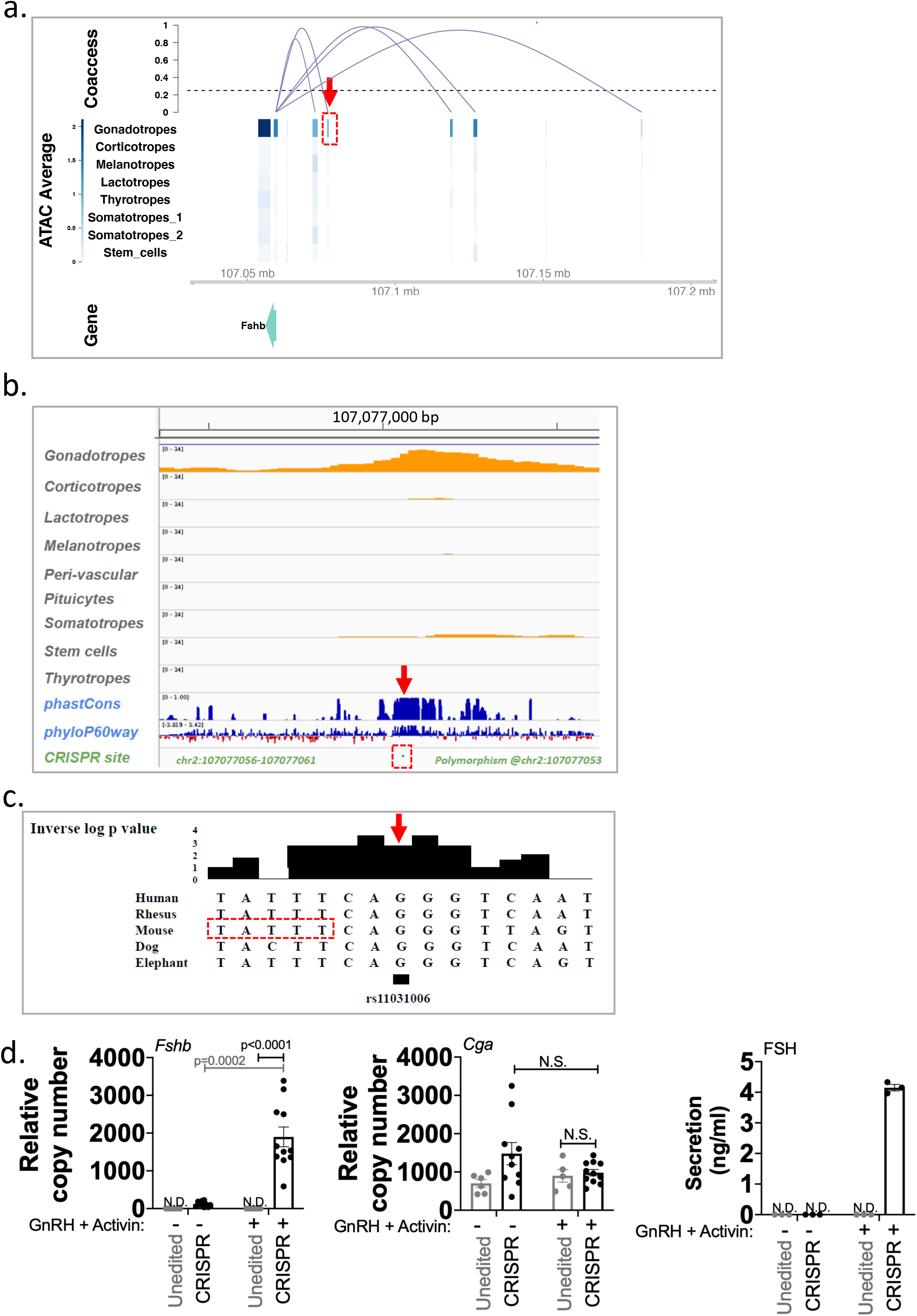
Genome-wide inference of pituitary cis-regulatory domains: *Fshb* gene case study. **a**. Cis-regulatory domains of the *Fshb* gene inferred from snATACseq data correlation in a single male pituitary (analysis of all 6 individual pituitaries is presented in **Supplementary Figure 16)**. Boxed in red is the 17kb upstream region corresponding to the human SNP rs11031006. **b**. Conservation analysis of the boxed region indicated in **a**. **c**. Multi-species sequence alignment of the *FSHB* SNP rs11031006 (arrow). The boxed sequence was deleted by CRISPR. **d**. Shown are *Fshb* (Left panel) and *Cga* (Center panel) gene expression, and FSH secretion levels (Right panel) in the unedited gonadotrope cell line (grey) and in the CRISPR deletion mutant line (black), either at baseline (−), or following GnRH and activin A treatment (+). Each data point is shown. Biological replicates were n=4 (*Fshb*), n=2 (*Cga*), each measured in technical triplicates, and n=3 (FSH). Error bars represent s.e.m. Significance was determined by ANOVA with Bonferroni corrections.

## DISCUSSION

We utilized an integrated, sn multi-omics analysis to elucidate the epigenetic mechanisms that regulate transcriptional networks in the pituitary. SnRNAseq and snATACseq assays were performed simultaneously on pituitaries from the same individual animals. Sn methylation profiling required the use of a pool of pituitaries to reach a sufficient number of nuclei for adequate genome-wide coverage. Our study identified epigenetically-defined cell type composition, sex-specific differences in transcription and epigenetic programs, an experimentally validated cis-regulatory domain, and epigenetic mechanisms contributing to cell type-specific regulon composition.

Using the PLIER framework, we identified transcriptional and accessibility LVs that varied by sex (**Fig. 6**, **Supplementary Figure 11**). Notably, distinctive gene expression and promoter accessibility patterns were found in females, and cell type-specific patterns were distinguished across individuals, revealing for instance putative differences in reproductive cycle stage. We identified corresponding transcriptional and accessibility LVs that varied in individual animals and propose that one of the females may have been non-cycling. Earlier studies identified a sexual difference in transcriptome-wide gene expression in the pituitary gland^42^, and recently in scRNAseq analysis of the mouse pituitary^3^. Similar to Ho et al., we observed a predominance of somatotropes in males compared to females, and a higher proportion of lactotropes in females relative to males. Understanding the sexual differences in the pituitary gland within both transcriptomic and epigenomic layers could have important clinical implications in humans, such as for the development of gender-specific treatments for pituitary disorders. While the present analysis is based on a small number of animals, it supports the value of capturing the *in vivo* expression and epigenetic state of each animal for simultaneous multi-omics assays. These results provide a foundation for future study of correlated transcriptional and epigenetic changes during the reproductive cycle and in response to perturbations at individual subject resolution.

Co-accessibility analysis provided a full genome compendium of putative cis-regulatory domains (see **Fig. 9a**, snpituitaryatlas.princeton.edu). One candidate was in a highly conserved co-accessible region 17 kb upstream of the *Fshb* gene that corresponds to the site of the human SNP rs11031006. While we were unable to experimentally target the SNP itself, CRISPR genetic ablation within this putative cis-regulatory locus in a gonadotrope-like cell line resulted in increased *Fshb* mRNA and FSH levels (**Fig. 9**). Similarly, increased *Fshb* mRNA levels were also observed with two other targeted deletions that were also in close proximity to the SNP. rs11031006 has been associated with alterations in female fertility, including endometriosis^39, 43^, longer menstrual cycles^44^, spontaneous dizygotic twinning^45^, and risk of polycystic ovary syndrome (PCOS^38^). While the presence of a sequence related to an estrogen response element has been noted in previous linkage studies, the mechanisms responsible for modulation of *Fshb* by the SNP or deletion by CRISPR remain to be determined. The effects of the CRISPR ablation on basal and stimulated *Fshb* expression support the value of the snATACseq co-accessibility resource to guide study of pituitary cis-regulatory domains.

RNA expression patterns alone cannot distinguish developmentally distinct cell types from transient and reversible alterations in cell states. Obtaining data from snap-frozen tissue eliminates the *ex vivo* alterations in expression patterns that may result from tissue handling, dissociation, or culturing ^46^. Analysis of our multi-omics sn data identifies and characterizes all the classical major pituitary cell types at the levels of RNA expression, chromatin accessibility and methylation (see **Figs. 2-5**, snpituitaryatlas.princeton.edu). Overall, cell type identification for all cell types was very similar in the snRNAseq and the snATACseq analysis, except for the detection of proliferating cells only in the transcriptomic analysis. Characterization of cell types based on the sn methylation data was comparable to that based on the other datasets for the primary pituitary cell types. However, sn methylation analysis identified fewer non-endocrine cell types and a homogeneous cluster of somatotropes. While sn methylation provides a deeper coverage per cell^23^, the number of nuclei assayed was considerably lower than in the other two assays, which may have contributed to discrepancies in cell type classification. However, methylation analysis has been considered the most reliable assay to distinguish cell subtypes^47^. Overall, we propose that identification of a specific cell lineage as opposed to a transient cell state requires confirmation by transcriptome, accessibility and methylation analysis. High cell number methylation studies will be needed to characterize the cell subtype clusters found by snRNAseq and snATACseq and determine whether they represent cell states or cell subtypes.

Analyses of the somatotrope cluster in the sn transcriptome and snATACseq data showed the presence of a gradually varying gradient of somatotropes. Gene ontology analysis using genes that were differentially expressed at the poles of this gradient showed an enrichment in cell adhesion, signaling, ion transport related genes in the somatotrope 1 pole and in genes involved in localization and cell junction organization in the somatotrope 2 pole. In males only, we identified a somatotrope 0 cell group closer to lactotropes which showed *Gh* but only minimal *Prl* expression (**Fig.2**). However, relatively lower nuclei numbers in the corresponding male snATACseq datasets make further discussion of these cells premature. Interestingly, earlier studies had described subpopulations of somatotropes in rat and pig^48, 49, 50^, and recent scRNAseq analysis of male murine pituitaries identified sub-populations within the somatotropes^1^. Our analysis raises the possibility that what has been identified as somatotrope subpopulations might represent a gradual continuum of varying transcriptome and chromatin accessibility patterns across the somatotrope lineage.

The characteristics and function of individual cells depends on the composition and activity of GRNs. A GRN is composed of regulons, each of which consists of a TF or upstream regulator along with its potential direct target genes. Notably, using SCENIC analysis for inferring pituitary regulons from snRNAseq data, we found that the pituitary cell types can be clustered and thus defined by their relative regulon activity levels (**Fig. 7a,b**). Thus, regulon activity may be considered a characteristic feature defining the physiology of each pituitary cell type. SCENIC identifies regulons based on correlated expression across all cell types; thus, genes that may not be expressed in a given cell type can be included in the set of regulon targets. Given the importance of FOXL2 in reproductive physiology^51, 52, 53^, we investigated the composition and control mechanisms of the FOXL2 regulon in gonadotropes. One cell type control mechanism for a regulon involves cell type-specific expression of the driver. *Foxl2* showed higher expression in gonadotropes than in other cell types (**Fig. 7d**). Approximately one-third of the FOXL2 regulon targets identified by SCENIC were presumed to be off in gonadotropes based on relative gene expression. Notably, we found a high correspondence between increased gonadotrope expression of FOXL2 regulon targets and gene accessibility (**Fig. 7e,f**). Yet, we found only a borderline negative association between gonadotrope expression and promoter methylation levels (**Supplementary Figure 12**). These results suggest that methylation state is only one factor contributing to the promoter accessibility state of potential regulon targets within a cell type. We demonstrate that a similar gene expression and chromatin accessibility correlation is found for other cell type-specific regulons, which influences their associated biological processes (**Fig. 8**, **Supplementary Figures 14, 15**). We propose that the composition of a cell type-specific regulon is determined by cell type-restricted-expression of the driver, the presence of binding sites on potential targets, and the epigenetic state of potential targets (**Fig. 8e**). Elucidating the interaction between the epigenetic mechanisms contributing to gene chromatin accessibility and cell type-specific regulon composition is an important question for further study.

The pituitary atlas resource is a comprehensive, integrated collection of sn transcriptome, chromatin accessibility, and DNA methylome datasets for the body’s “master gland”. This data integration has enabled us to classify cell types in the adult mouse pituitary, to dissect cellular heterogeneity within both transcriptomic and epigenomic layers, and to identify transcriptional regulatory mechanisms in specific cell populations. Our work lays the foundation for characterizing the epigenetic regulatory principles that control cell type-specific gene expression in the pituitary.

## METHODS

### Animals and pituitary collection

Pituitaries were collected from male and randomly-cycling C57BL/6 female mice aged 10-12 weeks. Animals were on a 12-hour on, 12-hour off light cycle (lights on at 7 AM; off at 7 PM). Pituitaries were immediately snap-frozen following dissection, and stored at −80C.

### Nuclei isolation from pituitaries

On ice, snap-frozen pituitaries were thawed and prepared based on a modified protocol from^54^. Briefly, RNAse inhibitor (NEB MO314L) was added to the homogenization buffer (0.32 M sucrose, 1 mM EDTA, 10 mM Tris-HCl, pH 7.4, 5mM CaCl2, 3mM Mg(Ac)2, 0.1% IGEPAL CA-630), 50% OptiPrep (Stock is 60% Media from Sigma; cat# D1556), 35% OptiPrep and 30% OptiPrep right before isolation. Each pituitary was homogenized in a dounce glass homogenizer (1ml, VWR cat# 71000-514), and the homogenate filtered through a 40 μm cell strainer. An equal volume of 50% OptiPrep was added, and the gradient centrifuged (SW41 rotor at 9200rpm; 4C; 25min). Nuclei were collected from the interphase, washed, resuspended either in 1X nuclei dilution buffer for snATACseq (10X Genomics) or in 1X PBS/0.04% BSA for snRNAseq, and counted (Invitrogen Countess II).

### SnRNAseq assay

SnRNAseq was performed following the Single Cell 3’ Reagents Kits V3 User Guidelines (10x Genomics, Pleasanton, CA). Nuclei were filtered and counted on a Countess instrument. A minimum of 1,000 nuclei were loaded (Chromium Single Cell 3’ Chip kit A v2 PN-12036 or v3 chip kit B PN-2000060). Reverse-transcription (RT) was performed in the emulsion, cDNA amplified, and library constructed with v2 or v3 chemistries. Libraries were indexed for multiplexing (Chromium i7 Multiplex kit PN-12062).

### SnRNAseq data analysis

SnRNAseq data were processed using the Cell Ranger pipeline v3.0.2, and aligned to whole transcripts rather than exonic regions only so as to include pre-mRNA. This resulted in a 2.15-fold increase in the median number of genes detected per cell in females and 2.3−2.4 fold increase in males relative to a traditional exonic-only alignment. Clustering and differential gene expression analysis were performed using Seurat v.3.1.1 and standard procedures^55, 56^. SnRNAseq data contain a background level of the most expressed genes in the sample. We first identified gel-beads in emulsion (GEMs) that contained only background as a peak in the UMI count distribution around 255 counts. We used troughs at both the high-end and low-end of the background peak to delimit the range of UMI counts to include in our background calling. Typically, this gave us a UMI counts range of roughly 150 to 400. GEMs that include cells were called with a minimum UMI count of 500 in Cell Ranger, and were thus well separated from those consisting of pure background. We verified that all expressed genes correlate linearly with the total UMI count within background gel-beads, meaning that their proportions did not change. Next, we added 200 random background barcodes into the output Cell Ranger cell-filtered feature matrices, and repeated the clustering. Background GEMs all clustered together and separated from any of the cell clusters (except for the debris cluster), confirming their homogeneous and distinct nature. We used our background identified GEMs to derive the background distribution of each expressed gene. While only a handful of transcripts were present at a concentration larger than one molecule per gel-bead, they could nevertheless reach counts of up to ~100 in the case of *Gh* in male samples and *Prl* in female samples.

### SnATACseq assay

SnATACseq was performed following the Chromium Single Cell ATAC Reagent Kits V1 User Guide (10x Genomics, Pleasanton, CA). Nuclei were counted (Countess counter), transposition was performed in 10μl at 37C for 60min on at least 1,000 nuclei, before loading of the Chromium Chip E (PN-2000121). Barcoding was performed in the emulsion (12 cycles) following the Chromium protocol. Libraries were indexed for multiplexing (Chromium i7 Sample Index N, Set A kit PN-3000262).

### SnATACseq analysis

SnATACseq data were processed using Cell Ranger-ATAC pipeline version 1.2.0 that eliminates barcode multiplets. We used the pipeline’s intrinsic *K-Means* clustering analysis to annotate cell types, as graph-based clustering tended to over-cluster. We employed a *k =* 10 (females), and a *k =* 7−9 (males), as they had fewer sequenced cells. Cell types were annotated based on cutsite pileup sums at promoter regions of known cell type markers. Cell types present in too low numbers for the *K-Means* algorithm (which is limited to k = 10) were identified as a separate cluster (for example non-pituitary cells). Nevertheless, we were able to visually locate these less abundant cell types as split on the tSNE plot and identify them using promoter sums of known markers. We identified doublets similarly. In male samples, which contain fewer nuclei, we were unable to split non-pituitary cells apart since their numbers were too low. Signac v. 0.1.5 was used to perform and confirm clustering on one male and one female sample using standard procedures.

### Quality control (QC) and sequencing of libraries

QC and quantification of libraries were done by Bioanalyzer (High-Sensitivity DNA Bioanalyzer kit) and mi-seq (Illumina). Sequencing was carried out at the New York Genome Center (NYGC) on an Illumina Novaseq using 98+26 paired-end reads.

### Sn methylation assay

The snmC-seq2 protocol was applied to individual nuclei isolated from a pool of 30 adult male pituitaries, following a protocol that included bisulfite conversion and subsequent preparation of the snmC-seq2 libraries as previously described^23, 57^. Sequencing was done at the Salk Institute on Illumina Novaseq 6000 using 150 bp paired-end reads.

### SnmC-seq2 mapping and preprocessing

A versatile mapping pipeline (<u>cemba-data.rtfd.io</u>) was implemented for all the sc methylome-based technologies developed by the Ecker Lab^23, 57, 58^. The main steps of this pipeline included: 1) demultiplexing FASTQ files into sc; 2) reads-level QC; 3) mapping; 4) BAM file processing and QC; 5) final molecular profile generation. Details of the five steps for snmC-seq2 were described previously^39^. We mapped all the reads onto the mouse mm10 genome. After mapping, we calculated the methyl-cytosine counts and total cytosine counts for two sets of genomic regions in each cell. The non-overlapping 100kb genomic bins of the mm10 genome (generated by “bedtools makewindows -w 100000”), which was used for clustering analysis and ANN model training, and the gene body region ± 2kb defined by the mouse GENCODE vm22, which was used for cluster annotation and integration with other modalities.

### Clustering analysis and DMR calling

Before clustering, we first filtered the cells based on these main mapping metrics: 1) mCCC rate < 0.03. mCCC rate reliably estimates the upper bound of bisulfite non-conversion rate^57^, 2) overall mCG rate > 0.5, 3) overall mCH rate < 0.2, 4) total final reads > 500,000, 5) bismark mapping rate > 0.5. The clustering analysis was performed as previously described^39^. In brief, we used the mCG and mCH 100kb genomic bins matrices as inputs of the clustering analysis. After calculating the normalized posterior estimation of mCG and mCG rate based on the beta-binomial distribution, we selected the top 2500 highly variable features and performed Principal Component Analysis (PCA) on each matrix and concatenate the first 20 PCs from both matrices. We performed KNN-Louvain clustering (K=25, Louvain resolution = 0.8) on the concatenated PCs. We manually annotated clusters based on marker genes learned from other modalities. After clustering, we merged the sc ALLC files (whole-genome cytosine count table for each cell) by cluster to get pseudo-bulk ALLC files. We then performed differential methylated regions (DMR) analysis using the “methylpy DMRfind” function among all clusters, as previously described^59^.

### PLIER data analysis

To examine more deeply the trends in gene expression of assigned cell types across samples and data types, we treated each sc dataset as a collection of bulk datasets for given labeled cell types. Each cell type was then treated as a separate bulk measurement within each sample. For snATACseq data, peak counts for a given gene were generated by selecting the peak closest to the transcription start site (TSS). These peak counts per gene were then collected into single bulk measurements for each cell type in each sample. We focused specifically on eight relevant cell types in the pituitary: corticotropes, gonadotropes, lactotropes, melanotropes, proliferating cells, somatotropes, stem/progenitor cells, and thyrotropes. For the snRNAseq dataset, this process generated 48 bulk measurements over six samples (three females and three males), and for the snATACseq dataset, we generated 42 bulk measurements. Proliferating cells were not identified in snATACseq. We applied PLIER^24^, which finds patterns in count data that are associated with known prior information (such as Reactome and Kegg). PLIER was run on each set of samples separately with LVs generated on the bulk measurements in an unsupervised fashion. LVs were then curated to find patterns relevant to individual cell types as well as sample-wide trends such as sex-based differences. Statistical significance of LVs was computed through the Kruskal-Wallis non-parametric test for multiple groups as part of the stat_compare_means R method. Comparisons between LVs within and across datatypes were achieved by comparing the overlap of the 50 genes most associated with a given LV.

### Generation of the CRISPR LβT2 clone

The murine LβT2 cell line^60, 61^ was a gift from Dr. Pamela Mellon (UCSD). Cells were maintained in 10-cm diameter dishes containing Dulbecco’s Modified Eagle’s Medium (DMEM) supplemented with 10% fetal bovine serum (FBS) and 1% penicillin/streptomycin (Invitrogen, Carlsbad, CA) in a humidified incubator at 37°C and 5% CO_2_. A 20 bp target site GCCCTGTGATATTTAT*TT*CA was chosen to induce double-strand DNA breaks at the TT location using CRISPR-Cas9 (University of Utah Mutation Generation and Detection Core). LβT2 cells were transfected at 50% confluency with 4 μg CRISPR Cas9-gRNA-GFP constructs, and 1 μg of a gel-purified mutagenic primer targeting mouse rs11031006 (5’-CTGGAATTTAATATTGCTCTGCCCTGTGATATTTATTTCA<u>A</u>GGTTAGTAGAAATGTAGCTACCTCCTGTAATGACAAATGA-3’) using PolyJet In Vitro DNA Transfection Reagent (SignaGen Laboratories). At 18 hours post transfection, cells were washed with PBS, digested with 0.5 ml 0.25% trypsin-EDTA, and reconstituted with 0.5 ml DMEM. Cells were sorted for GFP expression and collected by an Avalon Cell Sorter (Propel Labs, Fort Collins, CO), before being plated on a 12-well plate in supplemented DMEM. Isolation, genotyping, and expansion of individual colonies were repeated three times to establish new clonal cell lines. The QIAamp DNA Mini Kit (QIAGEN, Hilden, Germany) was used to extract DNA of each colony for verification of the clonal genotype (rs11031006 forward: 5’-TGAATGCTATTTTGTGGCAACT-3’ and rs11031006 reverse: 5’-TCATTTGTCATTACAGGAGGTAGC-3’) by Sanger sequencing. Genotype was identified using Poly Peak Parser ^62^. One clone was selected for further studies.

### Quantitative real-time PCR (qPCR)

Edited and unedited LβT2 cells were plated at 1,500 - 5,500 cells per well into 96-well plates in DMEM with 10% fetal bovine serum and 1% penicillin/streptomycin. Cells were treated for 24 hours either with vehicle or with 10nM GnRH and 1nM activin A (R&D Systems, Minneapolis, MN) in serum-free DMEM with 0.1% BSA.

Total RNA from each well was extracted using the RNeasy Mini Kit (Qiagen, Valencia, CA) and reverse transcribed to cDNA using SuperScript VILO MasterMix (ThermoFisher, Rockford, IL). Samples were then subjected to qPCR using PowerUp SYBR Green Master Mix (ThermoFisher, Rockford, IL) and the following primer sets detecting three transcripts: *Fshb* forward: 5’-GAAGAGTGCCGTTTCTGCAT-3’, *Fshb* reverse: 5’-CCGAGCTGGGTCCTTATACA-3’, *Cga* (*α-subunit*) forward: 5’-AGCTAGGAGCCCCCATCTAC-3’, *Cga* (*α-subunit*) reverse: 5’-TTCTCCACTCTGGCATTTCC-3’, *Gapdh* forward 5’-GGCATTGCTCTCAATGACAA-3’, *Gapdh* reverse 5’-TGTGAGGGAGATGCTCAGTG-3’. qPCR reactions were carried out with the following conditions: 50°C for 2 min; 95°C for 10 min; 40 cycles of 95°C for 15 sec and 60°C for 1 min using an Applied Biosystems 7900HT (Foster City, CA). SDS 2.1 software was used to identify the cycle number for each target (Applied Biosystems, Carlsbad, CA). Three technical qPCR replicates were run for each biological replicate. Results were exported as cycle threshold (Ct) values, and Ct values of target genes were normalized to those of *Gapdh* in subsequent analysis. Data were expressed as arbitrary units by using the formula, *E* = 2,500 × 1.93(^*Gapdh* CT value − gene of interest CT value)^, where *E* is the expression level in arbitrary units. The data were expressed as the mean ± s.e.m, and a value of *p-value* < 0.05 was considered statistically significant.

### Radioimmunoassays

FSH secreted in the medium was assayed, and the reportable range for the FSH assay was 1.1 to 54.1 ng/mL. The intra-assay and inter-assay %CV for FSH were 7.4% and 9.1% respectively.

### Sn data integration

The snRNAseq, snATACseq and sn methylation data were integrated in a reference-query based manner, mainly using the “FindTransferAnchors” and “TransferData” functions from the Seurat v3 package^55, 56^. The snRNAseq datasets were used as the reference and the other modalities were integrated to them. To integrate snATACseq to snRNAseq, the peak-by-cell accessibility matrix was converted to a gene-by-cell activity matrix based on the chromatin accessibility within each gene’s gene body and a 2kb upstream region, under the assumption that chromatin accessibility and gene expression were positively correlated. The variable features from the snRNAseq data were used to find the anchors and the snATACseq data in the LSI low-dimensional embedding were used to transfer the data from snRNAseq to snATACseq. To integrate the sn methylation data to snRNAseq, the normalized methylation rates for each gene’s gene body and +/− 2kb flanking region were used. This gene-by-cell methylation rate matrix was converted to an activity matrix by “activity = 1 / normalized methylation rate”, under the assumption that methylation and gene expression are negatively correlated. The variable features from the snRNAseq data were used to find the anchors and the sn methylation data in the PCA low-dimensional embedding were used to transfer the data from snRNAseq to sn methylation. Co-embedding of the 3 modalities was done by concatenating snRNAseq with the transferred data of snATACseq and sn methylation, and running dimension reduction on the concatenated dataset.

### SCENIC analysis

We used SCENIC^32^ to reconstruct the GRNs from snRNAseq data, with 3 steps: (1) identifying sets of genes that are co-expressed with TFs in the snRNAseq dataset, (2) refining the target genes for each TF based on enrichment of its cis-regulatory binding motifs, and (3) measuring the enrichment of each regulon (consisting of a TF and its target genes) in each nucleus using an AUC score (regulon activity score). SCENIC was run on filtered and pooled datasets of 3 females and 3 males with the raw UMI counts used as an input. In each dataset, only genes with UMI count >= 3 in at least 3 cells were kept, and cells identified as doublets were removed.

The output was a regulon-by-cell matrix composed of the activity of each regulon in each cell type. Cell type-level regulon activities were calculated by the average of sn-level regulon activities. For a regulon in a cell type, the relationship between chromatin accessibility and RNA expression of target genes were determined using their relative levels, defined as the ratio between cell type average and whole-dataset average. Chromatin accessibility of the target genes was measured by the accessibility either in the promoter region (500bp region in the upstream of TSS) or in the gene body region. We determined the relationship between relative gene accessibility and expression by scatter plots and linear regression.

To study the target genes specifically turned on in each regulon and each cell type, we identified a target gene as specifically highly expressed in a cell type if it satisfied either of these two conditions: (1) its relative averaged normalized RNA expression in this cell type vs. the whole dataset was greater than 2, and it had non-zero UMI counts in at least 10% of the cells in this cell type; and (2) it had non-zero UMI counts in at least 50% of the cells, and its relative averaged normalized RNA expression in this cell type was greater than 1. In the analysis involving multiple samples such as all males, a target gene was defined as specifically highly expressed in a cell type if it was specifically highly expressed in the cell type in at least one sample.

### Co-accessibility and putative regulatory region

Co-accessibility between all pairs of snATACseq peaks within 500kb was calculated using the Cicero package^37^. To find putative distal regulatory regions for a gene, we used the snATACseq peaks within the gene promoter region, and identified their co-accessible peaks that were at least 5 kb away.

### Functional module analysis of target genes

To identify the functions enriched in the lists of on and off genes in each regulon, we used the functional module detection method from the HumanBase resource (https://hb.flatironinstitute.org/module). The method clusters genes by their connectivities in a tissue-specific functional network, and finds enriched GO terms for each of the gene clusters.

### Conservation analysis

PhastCon and PhyloP(60) analyses were performed to analyze conservation among vertebrate species^40, 41^. PhyloP was performed on 60 vertebrate species.

### Statistics

For the assessment of sex difference in cell type proportions (**Fig. 4a,b**), we used a two-way analysis of variance (ANOVA) followed by Bonferroni multiple comparisons post-hoc test, with n = 3 biological replicates per cell type, F(11, 47)= 20.80 for snRNAseq, and F(8, 15)= 23.77 for snATACseq.

For analysis of LV sex differences by PLIER (**Fig. 6a**), we used a two-tailed Wilcoxon rank-sum test with n=48 (8 cell types, 6 biological samples), 1 degree of freedom, W = 490, p = 1.108e-05 for the snRNAseq data (Left panel) and with n=12 (2 cell types, 6 biological samples), 1 degree of freedom, W = 36, p = 0.002165 for the lactotrope and somatotrope snATACseq data (Right panel). For evaluating the cell type differences of gonadotrope-specific LVs by PLIER (**Fig. 6b**), we used a two-tailed Kruskal-Wallis analysis to test whether all samples originated from the same distribution. For LV2rna applied to snRNAseq data (Left panel), n=48 (8 cell types, 6 biological samples), 7 degrees of freedom, χ^2^ = 34.561, p = 1.352e-05. For LV2atac applied to snATACseq data (Center panel), n = 42 (7 cell types, 6 biological samples), 6 degrees of freedom, χ^2^ = 37.34, p = 1.512e-06. For LV2rna applied to snATACseq data (Right panel), n = 42 (7 cell types, 6 biological samples), 6 degrees of freedom, χ^2^ = 30.813, p = 2.752e-05.

For all the boxplots comparing chromatin accessibility or methylation between highly and lowly expressed genes, two-sided Wilcoxon rank-sum tests (a.k.a. Mann-Whitney U tests) were used, and p-values were calculated by normal approximation. Specifically, the R core function wilcox.test was used. The number of genes tested and the W values from the Wilcoxon rank-sum tests were: 235 highly vs. 75 lowly expressed genes (W = 3388, **Fig. 7f**), and 669 highly vs. 957 lowly expressed genes (W = 158752, **Fig. 8b**).

For *Fshb* expression (**Fig. 9d**), statistical analyses were all performed using GraphPad Prism version 5.04 (GraphPad Software, San Diego, CA, www.graphpad.com). We used a two-way ANOVA followed by Bonferroni multiple comparisons post-hoc test, with n = 4 biological replicates per sample (each measured in technical triplicates), F(1, 29)= 29.54 in the GnRH + activin A treatment in the unedited vs. CRISPR clones, and F(1, 29)= 22 in the CRISPR clone for the basal vs. GnRH + activin A treatment.

### Ethical compliance

We have complied with all ethical regulations and institutional protocols. All murine work was conducted at McGill University (Montreal, Quebec, Canada) under animal use protocol 5204, as approved by the Facility Animal Care Committee of the Goodman Cancer Research Centre.

### Data availability

The datasets (scRNAseq, snRNAseq, snATACseq) and sn methylation data generated in the present study are deposited in GEO. The sn mouse pituitary multi-omics atlas can be browsed via a web-based portal accessible at snpituitaryatlas.princeton.edu.

### Code availability

Any computational code used in the paper is available upon request.

## Acknowledgements

This work was supported by funding from the National Institute of Health (NIH) Grant DK46943 (SCS), the Canadian Institutes of Health Research (CIHR) Project Grants PJT-162343 (DJB) and PJT-169184 (DJB), NIH award R01HD065029 from the Eunice Kennedy Shriver National Institute Of Child Health & Human Development (CKW), NIH NICHD R01HD093461 (GVC), NIH R01HD087057 (GVC), and NIH NIDDK 1R01DK113776-01 (GVC). We acknowledge the Mutation Generation and Detection Core for reagents (University of Utah), the Flow Cytometry Core (University of Utah), the Genomic Core (University of Utah), and the New York Genome Center for sequencing. Radioimmunoassays were performed at the University of Virginia Core Ligand and Assay Laboratory.

## Author contributions

FRZ designed and performed research, analyzed and interpreted data, drafted the manuscript; ZZ, MZ, GRS, and GN contributed analytic tools and analyzed data; HL performed research and analyzed data; MM, RGC, NM, VDN, NS, MAA, XZ, LO, GS, CT, JRN, AB, AA, and NJ performed research; HP drafted the manuscript; JLT analyzed and interpreted data; DJB analyzed and interpreted data and provided analytical tools; CKW designed and performed research, analyzed and interpreted data; OGT and JRE contributed analytic tools; GVC discussed the results; SCS conceived research, analyzed data, and drafted the manuscript. All authors edited the manuscript and approved its final version.

## REFERENCES

1. Cheung LYM, et al. Single-Cell RNA Sequencing Reveals Novel Markers of Male Pituitary Stem Cells and Hormone-Producing Cell Types. Endocrinology 159, 3910–3924 (2018).

2. Fletcher PA, et al. Cell Type- and Sex-Dependent Transcriptome Profiles of Rat Anterior Pituitary Cells. Front Endocrinol (Lausanne) 10, 623 (2019).

3. Ho Y, et al. Single-cell transcriptomic analysis of adult mouse pituitary reveals sexual dimorphism and physiologic demand-induced cellular plasticity. Protein Cell, (2020).

4. Ruf-Zamojski F, et al. Single-cell stabilization method identifies gonadotrope transcriptional dynamics and pituitary cell type heterogeneity. Nucleic Acids Res 46, 11370–11380 (2018).

5. Mayran A, et al. Pioneer and nonpioneer factor cooperation drives lineage specific chromatin opening. Nat Commun 10, 3807 (2019).

6. Ludwig CH, Bintu L. Mapping chromatin modifications at the single cell level. Development 146, (2019).

7. Miller JL, Grant PA. The role of DNA methylation and histone modifications in transcriptional regulation in humans. Subcell Biochem 61, 289–317 (2013).

8. Buenrostro JD, et al. Single-cell chromatin accessibility reveals principles of regulatory variation. Nature 523, 486–490 (2015).

9. Clark SJ, Lee HJ, Smallwood SA, Kelsey G, Reik W. Single-cell epigenomics: powerful new methods for understanding gene regulation and cell identity. Genome Biol 17, 72 (2016).

10. Shema E, Bernstein BE, Buenrostro JD. Single-cell and single-molecule epigenomics to uncover genome regulation at unprecedented resolution. Nat Genet 51, 19–25 (2019).

11. Bakken TE, et al. Single-nucleus and single-cell transcriptomes compared in matched cortical cell types. PLoS One 13, e0209648 (2018).

12. Wu H, Kirita Y, Donnelly EL, Humphreys BD. Advantages of Single-Nucleus over Single-Cell RNA Sequencing of Adult Kidney: Rare Cell Types and Novel Cell States Revealed in Fibrosis. J Am Soc Nephrol 30, 23–32 (2019).

13. Lacar B, et al. Nuclear RNA-seq of single neurons reveals molecular signatures of activation. Nat Commun 7, 11022 (2016).

14. van den Brink SC, et al. Single-cell sequencing reveals dissociation-induced gene expression in tissue subpopulations. Nat Methods 14, 935–936 (2017).

15. Nguyen QH, Pervolarakis N, Nee K, Kessenbrock K. Experimental Considerations for Single-Cell RNA Sequencing Approaches. Front Cell Dev Biol 6, 108 (2018).

16. Mereu E, et al. Benchmarking single-cell RNA-sequencing protocols for cell atlas projects. Nat Biotechnol, (2020).

17. Habib N, et al. Massively parallel single-nucleus RNA-seq with DroNc-seq. Nat Methods 14, 955–958 (2017).

18. Lake BB, et al. Neuronal subtypes and diversity revealed by single-nucleus RNA sequencing of the human brain. Science 352, 1586–1590 (2016).

19. Selewa A, et al. Systematic Comparison of High-throughput Single-Cell and Single-Nucleus Transcriptomes during Cardiomyocyte Differentiation. Sci Rep 10, 1535 (2020).

20. Cusanovich DA, et al. A Single-Cell Atlas of In Vivo Mammalian Chromatin Accessibility. Cell 174, 1309–1324 e1318 (2018).

21. Preissl S, et al. Single-nucleus analysis of accessible chromatin in developing mouse forebrain reveals cell-type-specific transcriptional regulation. Nat Neurosci 21, 432–439 (2018).

22. Rai V, et al. Single-cell ATAC-Seq in human pancreatic islets and deep learning upscaling of rare cells reveals cell-specific type 2 diabetes regulatory signatures. Mol Metab 32, 109–121 (2020).

23. Luo C, et al. Robust single-cell DNA methylome profiling with snmC-seq2. Nat Commun 9, 3824 (2018).

24. Mao W, Zaslavsky E, Hartmann BM, Sealfon SC, Chikina M. Pathway-level information extractor (PLIER) for gene expression data. Nat Methods 16, 607–610 (2019).

25. Alter O, Brown PO, Botstein D. Singular value decomposition for genome-wide expression data processing and modeling. Proc Natl Acad Sci U S A 97, 10101–10106 (2000).

26. Deschenes J, Bourdeau V, White JH, Mader S. Regulation of GREB1 transcription by estrogen receptor alpha through a multipartite enhancer spread over 20 kb of upstream flanking sequences. J Biol Chem 282, 17335–17339 (2007).

27. Ghosh MG, Thompson DA, Weigel RJ. PDZK1 and GREB1 are estrogen-regulated genes expressed in hormone-responsive breast cancer. Cancer Res 60, 6367–6375 (2000).

28. Hodgkinson K, Forrest LA, Vuong N, Garson K, Djordjevic B, Vanderhyden BC. GREB1 is an estrogen receptor-regulated tumour promoter that is frequently expressed in ovarian cancer. Oncogene 37, 5873–5886 (2018).

29. Mohammed H, et al. Endogenous purification reveals GREB1 as a key estrogen receptor regulatory factor. Cell Rep 3, 342–349 (2013).

30. Speert DB, Sj MC, Seasholtz AF. Sexually dimorphic expression of corticotropin-releasing hormone-binding protein in the mouse pituitary. Endocrinology 143, 4730–4741 (2002).

31. Ma A, et al. IRIS3: integrated cell-type-specific regulon inference server from single-cell RNA-Seq. Nucleic Acids Res, (2020).

32. Aibar S, et al. SCENIC: single-cell regulatory network inference and clustering. Nat Methods 14, 1083–1086 (2017).

33. Ingraham HA, et al. The nuclear receptor steroidogenic factor 1 acts at multiple levels of the reproductive axis. Genes Dev 8, 2302–2312 (1994).

34. Stallings CE, Kapali J, Ellsworth BS. Mouse Models of Gonadotrope Development. Prog Mol Biol Transl Sci 143, 1–48 (2016).

35. Li Y, Schang G, Boehm U, Deng CX, Graff J, Bernard DJ. SMAD3 Regulates Follicle-stimulating Hormone Synthesis by Pituitary Gonadotrope Cells in Vivo. J Biol Chem 292, 2301–2314 (2017).

36. Becht E, et al. Dimensionality reduction for visualizing single-cell data using UMAP. Nat Biotechnol, (2018).

37. Pliner HA, et al. Cicero Predicts cis-Regulatory DNA Interactions from Single-Cell Chromatin Accessibility Data. Mol Cell 71, 858–871 e858 (2018).

38. Day FR, et al. Causal mechanisms and balancing selection inferred from genetic associations with polycystic ovary syndrome. Nat Commun 6, 8464 (2015).

39. Matalliotakis M, et al. The role of gene polymorphisms in endometriosis. Mol Med Rep 16, 5881–5886 (2017).

40. Pollard KS, Hubisz MJ, Rosenbloom KR, Siepel A. Detection of nonneutral substitution rates on mammalian phylogenies. Genome Res 20, 110–121 (2010).

41. Siepel A, et al. Evolutionarily conserved elements in vertebrate, insect, worm, and yeast genomes. Genome Res 15, 1034–1050 (2005).

42. Nishida Y, Yoshioka M, St-Amand J. Sexually dimorphic gene expression in the hypothalamus, pituitary gland, and cortex. Genomics 85, 679–687 (2005).

43. Sapkota Y, et al. Meta-analysis identifies five novel loci associated with endometriosis highlighting key genes involved in hormone metabolism. Nat Commun 8, 15539 (2017).

44. Laisk T, et al. Large-scale meta-analysis highlights the hypothalamic-pituitary-gonadal axis in the genetic regulation of menstrual cycle length. Hum Mol Genet 27, 4323–4332 (2018).

45. Mbarek H, Dolan CV, Boomsma DI. Two SNPs Associated With Spontaneous Dizygotic Twinning: Effect Sizes and How We Communicate Them. Twin Res Hum Genet 19, 418–421 (2016).

46. Denisenko E, et al. Systematic assessment of tissue dissociation and storage biases in single-cell and single-nucleus RNA-seq workflows. Genome Biol 21, 130 (2020).

47. Lee DS, et al. Simultaneous profiling of 3D genome structure and DNA methylation in single human cells. Nat Methods 16, 999–1006 (2019).

48. Hoeffler JP, Hicks SA, Frawley LS. Existence of somatotrope subpopulations which are differentially responsive to insulin-like growth factor I and somatostatin. Endocrinology 120, 1936–1941 (1987).

49. Castano JP, et al. Somatostatin increases growth hormone (GH) secretion in a subpopulation of porcine somatotropes: evidence for functional and morphological heterogeneity among porcine GH-producing cells. Endocrinology 137, 129–136 (1996).

50. Dobado-Berrios PM, et al. Heterogeneity of growth hormone (GH)-producing cells in aging male rats: in vitro GH releasing activity of somatotrope subpopulations. Mol Cell Endocrinol 123, 127–137 (1996).

51. Justice NJ, Blount AL, Pelosi E, Schlessinger D, Vale W, Bilezikjian LM. Impaired FSHbeta expression in the pituitaries of Foxl2 mutant animals. Mol Endocrinol 25, 1404–1415 (2011).

52. Tran S, et al. Impaired fertility and FSH synthesis in gonadotrope-specific Foxl2 knockout mice. Mol Endocrinol 27, 407–421 (2013).

53. Fortin J, Boehm U, Deng CX, Treier M, Bernard DJ. Follicle-stimulating hormone synthesis and fertility depend on SMAD4 and FOXL2. FASEB J 28, 3396–3410 (2014).

54. Mathys H, et al. Single-cell transcriptomic analysis of Alzheimer's disease. Nature 570, 332–337 (2019).

55. Butler A, Hoffman P, Smibert P, Papalexi E, Satija R. Integrating single-cell transcriptomic data across different conditions, technologies, and species. Nat Biotechnol 36, 411–420 (2018).

56. Stuart T, et al. Comprehensive Integration of Single-Cell Data. Cell 177, 1888–1902 e1821 (2019).

57. Luo C, et al. Single-cell methylomes identify neuronal subtypes and regulatory elements in mammalian cortex. Science 357, 600–604 (2017).

58. Luo C, et al. Single nucleus multi-omics links human cortical cell regulatory genome diversity to disease risk variants. Biorxiv https://doi.org/10.1101/2019.12.11.873398, (2019).

59. Schultz MD, et al. Human body epigenome maps reveal noncanonical DNA methylation variation. Nature 523, 212–216 (2015).

60. Graham KE, Nusser KD, Low MJ. LbetaT2 gonadotroph cells secrete follicle stimulating hormone (FSH) in response to active A. J Endocrinol 162, R1–5 (1999).

61. Turgeon JL, Kimura Y, Waring DW, Mellon PL. Steroid and pulsatile gonadotropin-releasing hormone (GnRH) regulation of luteinizing hormone and GnRH receptor in a novel gonadotrope cell line. Mol Endocrinol 10, 439–450 (1996).

62. Hill JT, Demarest BL, Bisgrove BW, Su YC, Smith M, Yost HJ. Poly peak parser: Method and software for identification of unknown indels using sanger sequencing of polymerase chain reaction products. Dev Dyn 243, 1632–1636 (2014).

