## Supplementary Information for "Single nucleus multi-omics regulatory atlas of the murine pituitary" (from another version of this preprint: version 1, 2020-06-06)

### 1. SUPPLEMENTARY METHODS

#### Animals and dissociation of primary murine pituitaries

*Standard protocol used for dissociation*

Pituitaries were collected from randomly cycling male and female C57BL/6 mice aged 10-12 weeks. Animals were on a 12-hour on, 12-hour off light cycle, with lights on at 7 AM and off at 7 PM. Upon collection, pituitaries were digested with collagenase and dispersed with pancreatin and the dissociated cells were collected acutely. About 0.7-2.5x10^6 cells were collected from single pituitaries and placed in individual tubes for cryopreservation. All mouse work was conducted at McGill University (Montreal, Quebec, Canada) under animal use protocol 5204, as approved by the Facility Animal Care Committee of the Goodman Cancer Research Centre.

*Improved protocol used for gentle dissociation with downstream SC assays:*

Upon removal, pituitaries were cut in half, rinsed 2x in Ca/Mg-free PBS and transferred to a tube containing trypsin solution (final concentration at 0.37 mg/ml, Sigma T9935) and gently agitated at 37°C for 10 minutes. After gentle aspiration and expulsion of the tissue, the removed supernatant was diluted with MEM/10% FBS and the process repeated for three cycles. The combined supernatants were centrifuged and the pellet resuspended in MEM/10%FBS. Approximately 1.7-2.5x10^6 cells were collected from single pituitaries.

**Cryopreservation/thawing of dissociated pituitary cells**

Dissociated pituitary cells were cryopreserved in 90% FBS/10% DMSO and shipped to New York on dry-ice. Upon arrival, cells were stored in liquid nitrogen. On the day of an experiment, the vial was removed from liquid nitrogen on dry-ice, thawed quickly in a 37C water bath, and cells were serially diluted with 10% FBS /DMEM until a minimum volume of 8ml was reached. Cells were centrifuged at 300xg for 5min, resuspended in 1ml of medium and counted.

**ScRNAseq assay**

ScRNAseq was performed as described (10x Genomics, Pleasanton, CA; (24)), following the Single Cell 3' Reagents Kits V2 or V3 User Guidelines. Cells were filtered, counted on a Countess instrument, and resuspended at a concentration of 1,000 cells/μl. The number of cells loaded on the chip was determined based on the 10X Genomics protocol. The 10X chip (Chromium Single Cell 3' Chip kit A v2 PN-12036 or v3 chip kit B PN-2000060) was loaded to target 5,000-10,000 cells final. Reverse-transcription was performed in the emulsion and cDNA was amplified before library construction following the Chromium protocols v2 or v3. Each library was tagged with a different index for multiplexing (Chromium i7 Multiplex kit PN-12062). Quality control and quantification of the amplified cDNA were assessed on a Bioanalyzer (High-Sensitivity DNA Bioanalyzer kit). Sequencing was carried out at the NYGC on Illumina Novaseq using 98+26 paired-end reads.

**Nuclei isolation from dissociated pituitary cells**

Nuclei were isolated from all cells remaining after the scGEM Drop-seq RNA-seq assay was done. Cells were centrifuged at 500g for 5min at 4C. The supernatant was carefully removed and 0.1X lysis buffer (1x: 10mM Tris-HCl pH 7.5, 10mM NaCl, 3mM $MgCl_2$, nuclease-free $H_2O$, 0.1% v/v NP-40, 0.1% v/v Tween-20, 0.01% v/v digitonin) was added. After 5min incubation on ice, 1ml of wash buffer was added. The nuclei were pelleted at 500xg for 5min at RT, resuspended either in 1X PBS/0.04% BSA for scGEM Drop-seq RNA-seq or in diluted nuclei buffer for snATAC-seq. Nuclei were counted and the concentration adjusted to run the given assay.

**PLIER analysis**

PLIER analysis was performed on scRNAseq studies done on dissociated pituitary cells from nine samples (four females and five males) applying the LV obtained from the snRNAseq data from the snap-frozen samples.

**Statistics**

For determining whether the association of PLIER LVs to known prior knowledge gene pathway information was random (**Supplementary Figure 10**), we applied a cross-validation test[1] that determines if each association between LV and prior knowledge pathway was robust to removing random one-fifth subsets of the genes associated with each pathway. AUC values were calculated based upon the ability to recover each pathway for a given LV.

For analysis of LV sex differences by PLIER applied to the dissociated pituitary dataset (**Supplementary Figure 11**), we used a two-tailed Wilcoxon rank-sum test with n=48 (8 cell types over 6 biological samples), 1 degree of freedom, W = 490, p = 1.108e-05 for the snap-frozen snRNAseq data (Left panel – as seen in **Fig. 6a**) and with n=64 (up to 8 cell types over 9 biological samples), 1 degree of freedom, W = 905, p = 1.142e-08 for the dissociated pituitary RNAseq data (Right panel).

For all the boxplots comparing chromatin accessibility or methylation between highly expressed and lowly expressed genes, two-sided Wilcoxon rank-sum tests (a.k.a Mann-Whitney U tests) were used, and p values were calculated by normal approximation. Specifically, the R core function wilcox.test was used. The numbers of genes tested and the W values from the Wilcoxon rank-sum tests were: in **Supplementary Figure 12a** compared 232 highly expressed vs. 75 lowly expressed genes (W = 8995); in **Supplementary Figure 12b** compared 25 highly expressed vs. 25 lowly expressed genes (W = 175); in **Supplementary Figure 14a** compared 284 highly expressed vs. 112 lowly expressed genes for males (W = 6771), and compared 332 highly expressed vs. 64 lowly expressed genes for females (W = 4419); in **Supplementary Figure 15b** compared 871 highly expressed vs. 755 lowly expressed genes (W = 182337).

**Data availability**

The datasets for the dissociated pituitary cells (scRNAseq, snRNAseq, snATACseq data) generated in the present study for dissociated pituitaries are deposited in GEO.

#### 2. SUPPLEMENTARY REFERENCES


1. Mao W, Zaslavsky E, Hartmann BM, Sealfon SC, Chikina M. Pathway-level information extractor (PLIER) for gene expression data. *Nat Methods* **16**, 607-610 (2019).

#### 3. SUPPLEMENTARY TABLES

##### Supplementary Table 1: Summary sample table

| Sample # | Sample name | Sinai ID | Species | Gender | Dissociation type | Material used | Assay | Estimated number of cells (cell Ranger) |
|---|---|---|---|---|---|---|---|---|
| 1 | Male mouse pit #2 | MM_Pit_2v2 | Mouse | Male | standard | cells | SC RNAseq v2 | 1612 |
| 2 | Male mouse pit #1 | MM_Pit_1v2 | Mouse | Male | standard | cells | SC RNAseq v2 | 542 |
| 3 | Male mouse pit #5 | MM_Pit_5 | Mouse | Male | standard | cells | SC RNAseq v3 | 3037 |
| 4 | Male mouse pit #6 | MM_Pit_6 | Mouse | Male | standard | cells | SC RNAseq v3 | 2476 |
| 5 | Female mouse pit #1 | FM_Pit_1-1 | Mouse | Female | gentle | cells | SC RNAseq v3 | 11974 |
| 6 | Female mouse pit #1 | FM_Pit_1-2 | Mouse | Female | gentle | nuclei | SN RNAseq v3 | 28109 |
| 7 | Male mouse pit #4 | MM_Pit_4-1 | Mouse | Male | standard | cells | SC RNAseq v3 | 2565 |
| 8 | Male mouse pit #4 | MM_Pit_4-2 | Mouse | Male | standard | nuclei | SN RNAseq v3 | 9901 |
| 9 | Female mouse pit #2 | FM_Pit_2-1 | Mouse | Female | gentle | cells | SC RNAseq v3 | 9674 |
| 10 | Female mouse pit #2 | FM_Pit_2-2 | Mouse | Female | gentle | nuclei | SN RNAseq v3 | 10162 |
| 11 | Female mouse pit #1 | Female_pit_1 | Mouse | Female | gentle | nuclei | SC ATAC, v1 | 4060 |
| 12 | Male mouse pit #4 | Male_pit_4 | Mouse | Male | standard | nuclei | SC ATAC, v1 | 20126 |
| 13 | Female mouse pit #2 | Female_Pit_2 | Mouse | Female | gentle | nuclei | SC ATAC, v1 | 7782 |
| 14 | Male pit#1 | froz_pit_MM1_R | Mouse | Male | none, snap-frozen | nuclei | SN RNAseq v3 | 7235 |
| 15 | Male pit#1 | froz_pit_MM1_A | Mouse | Male | none, snap-frozen | nuclei | SC ATAC, v1 | 2840 |
| 16 | Male pit#2 | froz_pit_MM2_R | Mouse | Male | none, snap-frozen | nuclei | SN RNAseq v3 | 4656 |
| 17 | Male pit#2 | froz_pit_MM2_A | Mouse | Male | none, snap-frozen | nuclei | SC ATAC, v1 | 967 |
| 18 | Male pit#3 | froz_pit_MM3_R | Mouse | Male | none, snap-frozen | nuclei | SN RNAseq v3 | 6215 |
| 19 | Male pit#3 | froz_pit_MM3_A | Mouse | Male | none, snap-frozen | nuclei | SC ATAC, v1 | 1082 |
| 20 | Female pit#1 | froz_pit_FM1_R | Mouse | Female | none, snap-frozen | nuclei | SN RNAseq v3 | 7383 |
| 21 | Female pit#1 | froz_pit_FM1_A | Mouse | Female | none, snap-frozen | nuclei | SC ATAC, v1 | 4968 |
| 22 | Female pit#2 | froz_pit_FM2_R | Mouse | Female | none, snap-frozen | nuclei | SN RNAseq v3 | 5744 |
| 23 | Female pit#2 | froz_pit_FM2_A | Mouse | Female | none, snap-frozen | nuclei | SC ATAC, v1 | 10248 |
| 24 | Female pit#3 | froz_pit_FM3_R2 | Mouse | Female | none, snap-frozen | nuclei | SN RNAseq v3 | 4474 |
| 25 | Female pit#3 | froz_pit_FM3_A | Mouse | Female | none, snap-frozen | nuclei | SC ATAC, v1 | 13338 |
| 26 | 30 pooled male pituitaries | SC methylation | Mouse | Males | none, snap-frozen | nuclei | SnmC seq2 | 2756 |

| | | | | |
|---|---|---|---|---|
| **Total cells SC RNAseq** | **31880** | cells | from 7 individual animals (5 males and 2 females) | 183926 |
| **Total cells SN RNAseq** | **83879** | nuclei | from 9 individual animals (5 females and 4 males) | |
| **Total cells ATAC seq** | **65411** | nuclei | from 9 individual animals (5 females and 4 males) | |
| **Total cells methylation** | **2756** | nuclei | from 30 pooled animals (all males) | |

Table showing all the samples run in the study, along with the dissociation and type of assay performed. The number of cells/nuclei sequenced for either sc or sn assays is also provided.

#### Supplementary Table 2: Cell type proportions

a.

| | FEMALES | | | | | MALES | | | | | Males/females |
|---|---|---|---|---|---|---|---|---|---|---|---|
| **Cell types** | **#1** | **#2** | **#3** | **Average** | **Std dev** | **#1** | **#2** | **#3** | **Average** | **Std dev** | **Stat. sign.** |
| lactotropes (%) | 31.22 | 38.72 | 37.67 | **35.87** | 4.06 | 20.91 | 15.08 | 19.30 | **18.43** | 3.01 | **Yes** |
| somatotropes (%) | 27.32 | 27.72 | 36.98 | **30.67** | 5.47 | 46.31 | 56.97 | 47.17 | **50.15** | 5.92 | **Yes** |
| melanotropes (%) | 9.11 | 7.06 | 5.57 | **7.25** | 1.78 | 8.78 | 5.85 | 8.80 | **7.81** | 1.70 | **No** |
| gonadotropes (%) | 3.20 | 2.84 | 3.80 | **3.28** | 0.48 | 6.55 | 5.41 | 6.41 | **6.12** | 0.62 | **No** |
| corticotropes (%) | 3.48 | 3.22 | 2.03 | **2.91** | 0.77 | 4.06 | 1.95 | 3.68 | **3.23** | 1.12 | **No** |
| thyrotropes (%) | 1.95 | 1.38 | 1.67 | **1.67** | 0.28 | 1.47 | 1.42 | 1.74 | **1.54** | 0.17 | **No** |
| stem cells (%) | 12.79 | 10.22 | 8.29 | **10.43** | 2.26 | 7.33 | 8.66 | 7.92 | **7.97** | 0.67 | **No** |
| proliferating (%) | 6.18 | 5.04 | 2.48 | **4.57** | 1.89 | 1.13 | 0.55 | 1.48 | **1.05** | 0.47 | **No** |
| Pericytes (%) | 1.64 | 1.74 | 0.45 | **1.28** | 0.72 | 1.06 | 1.29 | 1.37 | **1.24** | 0.16 | **No** |
| endothelial (%) | 0.89 | 1.04 | 0.00 | **0.64** | 0.56 | 0.80 | 1.21 | 1.03 | **1.01** | 0.20 | **No** |
| macrophages (%) | 0.64 | 0.55 | 0.36 | **0.52** | 0.14 | 0.66 | 0.46 | 0.36 | **0.49** | 0.15 | **No** |
| pituicytes (%) | | 0.47 | 0.69 | **0.58** | 0.16 | 0.95 | 1.14 | 0.75 | **0.95** | 0.20 | **No** |

b.

| | FEMALES | | | | | MALES | | | | | Males/females |
|---|---|---|---|---|---|---|---|---|---|---|---|
| **Cell types** | **#1** | **#2** | **#3** | **Average** | **Std dev** | **#1** | **#2** | **#3** | **Average** | **Std dev** | **Stat. sign.** |
| lactotropes (%) | 33.18 | 40.31 | 39.14 | **37.54** | 3.83 | 23.58 | 22.61 | 17.63 | **21.27** | 3.20 | **Yes** |
| somatotropes (%) | 25.54 | 27.92 | 32.30 | **28.59** | 3.43 | 43.21 | 56.48 | 48.22 | **49.30** | 6.70 | **Yes** |
| melanotropes (%) | 7.30 | 5.99 | 6.35 | **6.55** | 0.68 | 7.44 | 5.00 | 8.22 | **6.89** | 1.68 | **No** |
| gonadotropes (%) | 3.67 | 3.85 | 4.48 | **4.00** | 0.42 | 5.86 | 5.00 | 6.03 | **5.63** | 0.55 | **No** |
| corticotropes (%) | 5.12 | 4.11 | 4.82 | **4.68** | 0.52 | 4.10 | 3.52 | 3.38 | **3.67** | 0.38 | **No** |
| thyrotropes (%) | 2.50 | 1.79 | 2.59 | **2.29** | 0.43 | 1.62 | 1.14 | 1.19 | **1.31** | 0.26 | **No** |
| stem cells (%) | 10.40 | 8.70 | 9.27 | **9.46** | 0.86 | 6.90 | 6.25 | 6.85 | **6.67** | 0.36 | **No** |
| pituicytes (%) | 1.16 | 0.58 | 1.06 | **0.93** | 0.31 | 0.75 | | 3.74 | **2.25** | 2.11 | **No** |
| multiplets (%) | 11.14 | 6.75 | | **8.94** | 3.11 | 6.54 | | 4.75 | **5.65** | 1.27 | **No** |

c.

| | MALES |
|---|---|
| **Cell types** | Pooled |
| lactotropes (%) | 20.57 |
| somatotropes (%) | 41.80 |
| melanotropes (%) | 7.44 |
| gonadotropes (%) | 4.75 |
| corticotropes (%) | 4.17 |
| thyrotropes (%) | 2.21 |
| stem cells (%) | 8.13 |
| pericytes (%) | 3.16 |
| endithelial (%) | 6.39 |
| outliers (%) | 1.38 |

Table showing the percentages of cells obtained after analysis by snRNAseq (**a**.), snATACseq (**b**.), and sn-methylation (**c**.).

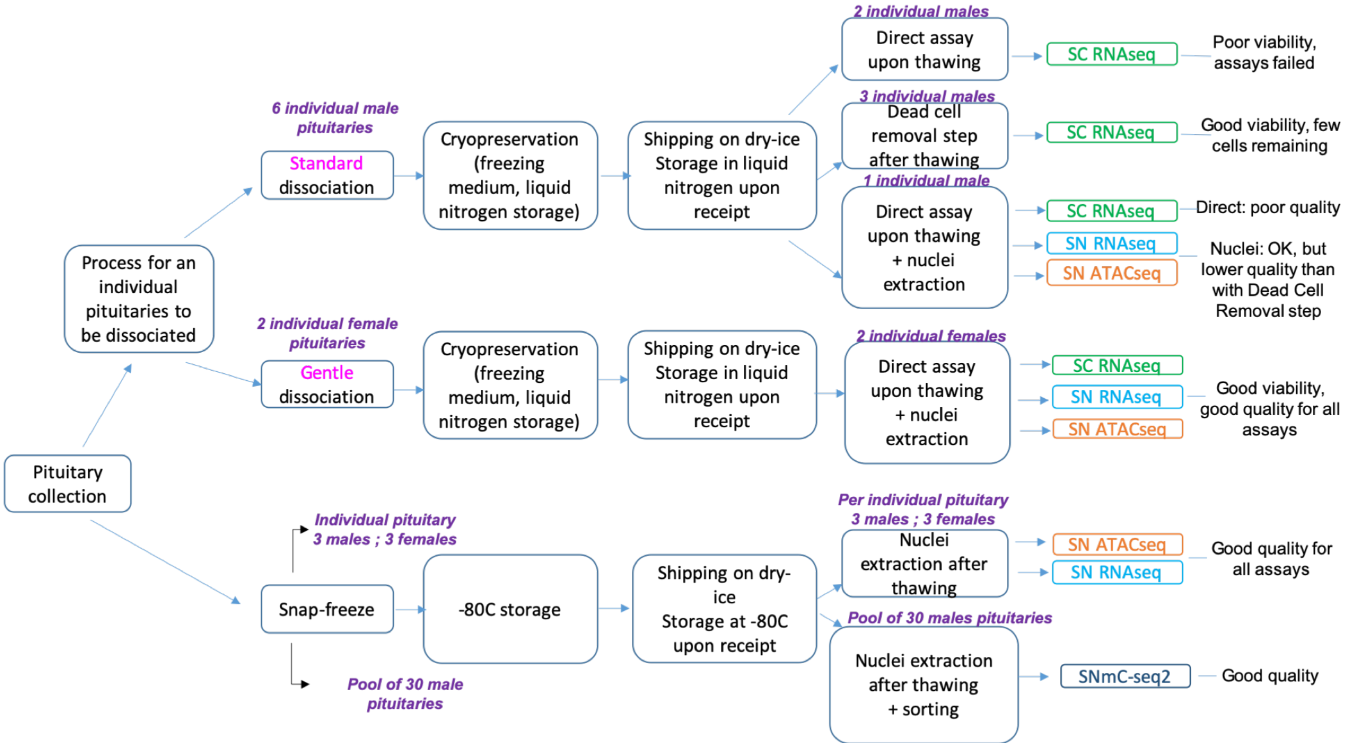


**Supplementary Figure 1:**
Schematic showing the different experimental strategies that were tested, from dissociated cells to whole snap-frozen pituitaries, standard to gentle cell dispersion, and the sc assays that were performed and their outcomes.

a.

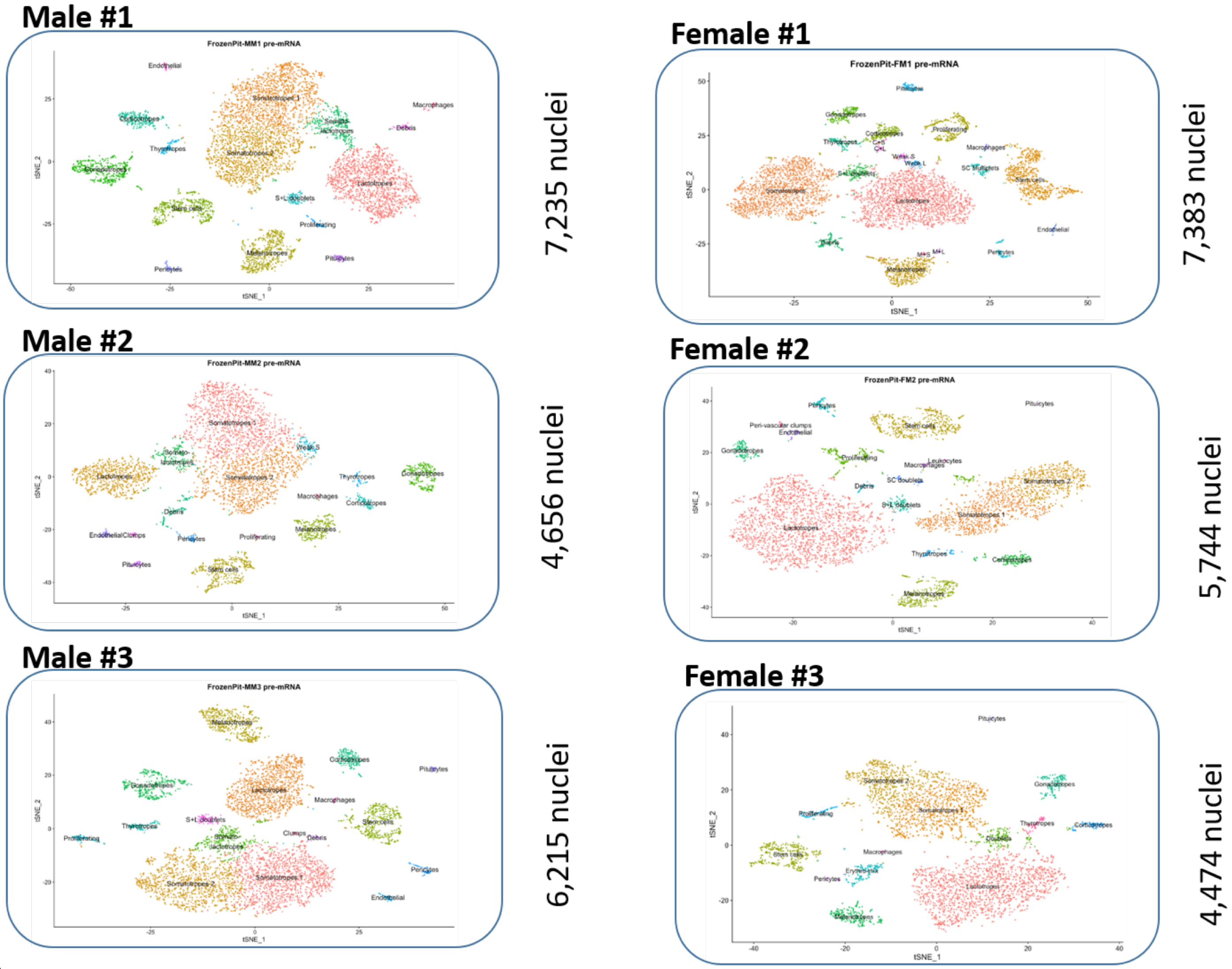


b.

| | FM1 | FM2 | FM3 | MM1 | MM2 | MM3 | Average |
|---|---|---|---|---|---|---|---|
| Number of cells | 7,406 | 5,725 | 4,515 | 7,322 | 4,726 | 6,279 | 5,996 |
| Mean reads/cell | 26,738 | 37,757 | 45,381 | 29,171 | 43,339 | 34,301 | 36,115 |
| Median genes/cell | 1,934 | 2,374 | 1,886 | 1,791 | 1,903 | 1,784 | 1,945 |
| Number of reads | 198,024,328 | 216,159,707 | 204,894,708 | 213,593,436 | 204,818,325 | 215,374,733 | 208,810,873 |
| Valid barcodes | 96% | 95.90% | 96.22% | 95.30% | 95.60% | 95.60% | 95.77% |
| Valid UMI | 100% | 100% | 100% | 100% | 99.90% | 99.90% | 99.97% |
| Sequencing saturation | 39.60% | 49.20% | 67.30% | 55.60% | 67% | 66.50% | 57.53% |
| Reads map to genome | 89.80% | 90.50% | 93.10% | 93.20% | 94% | 94.20% | 92.47% |
| Reads map to transcriptome | 46.50% | 48.50% | 59.10% | 50.00% | 52% | 39.20% | 49.23% |
| Total genes detected | 23,958 | 23,982 | 22,940 | 23,684 | 23,348 | 23,431 | 23,557 |

**Supplementary Figure 2:**
Results obtained for the snap-frozen pituitaries that were processed for snRNAseq. **a**. tSNE of all individual animals processed for snRNAseq from individually snap-frozen pituitaries (3 males and 3 females). **b**. Metrics obtained in all 6 samples. Refer to **Supplementary Fig. 7** for the same samples analyzed by snATACseq.

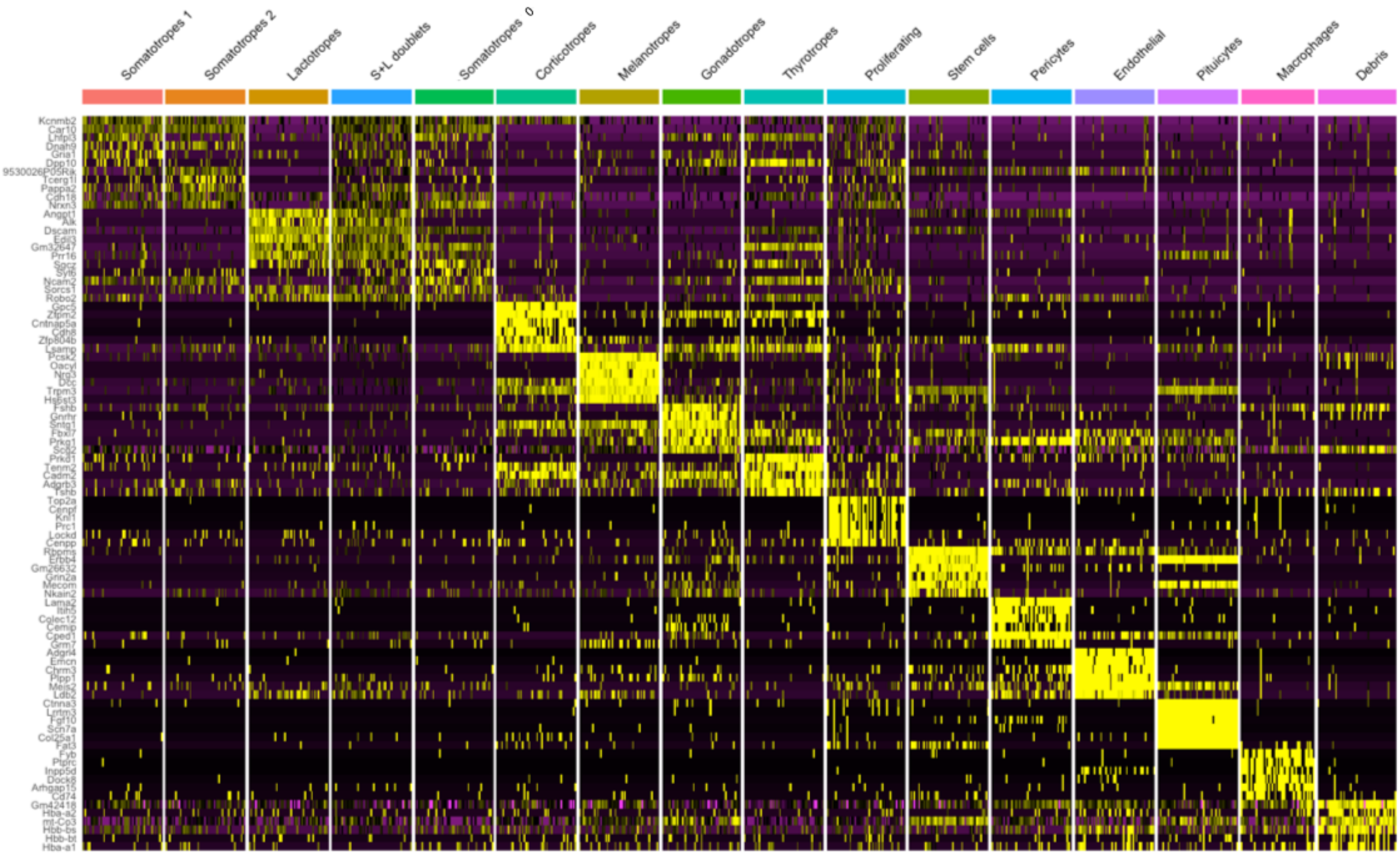


**Supplementary Figure 3:**
Heatmap showing cell identity based on gene expression in one male snap-frozen male pituitary.

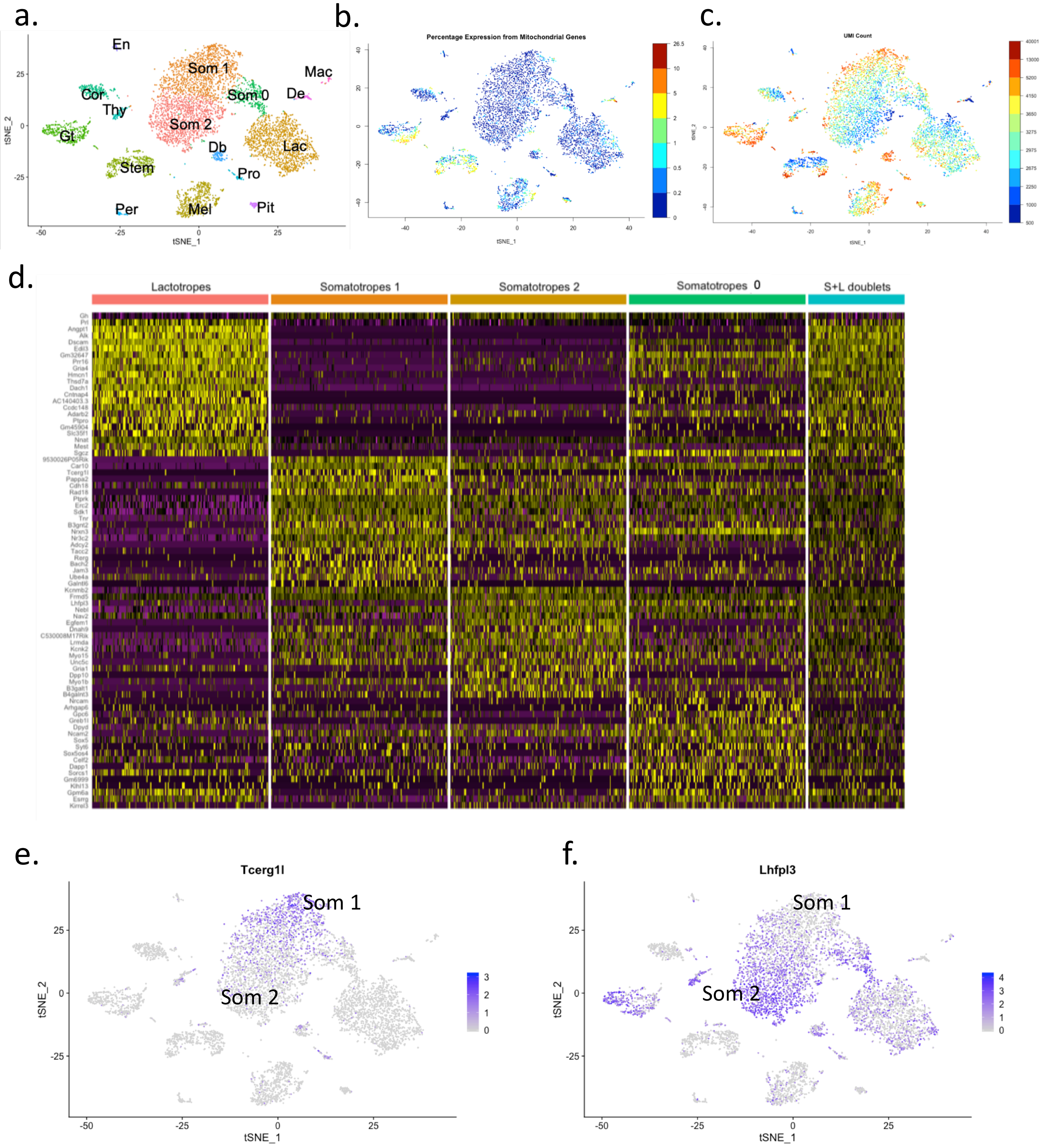


**Supplementary Figure 4:**
Analysis of the somatotrope gradient cluster and the link between somatotropes and lactotropes via somatotrope 0 (som 0). **a**. tSNE of the pituitary clusters. **b**. Mitochondrial analysis of the clusters showing a low mitochondrial content in the somatotrope and lactotrope clusters. **c**. UMI counts of the different clusters. **d**. Heatmap of the lactotropes, somatotrope sub-types, and doublets showing the differences in gene expression. **e, f.** Feature plots showing *Tcerg1l* expression at the Som 1 pole of the somatotrope cluster (**e**), and *Lhfpl3* expression at the Som 2 pole (**f**).

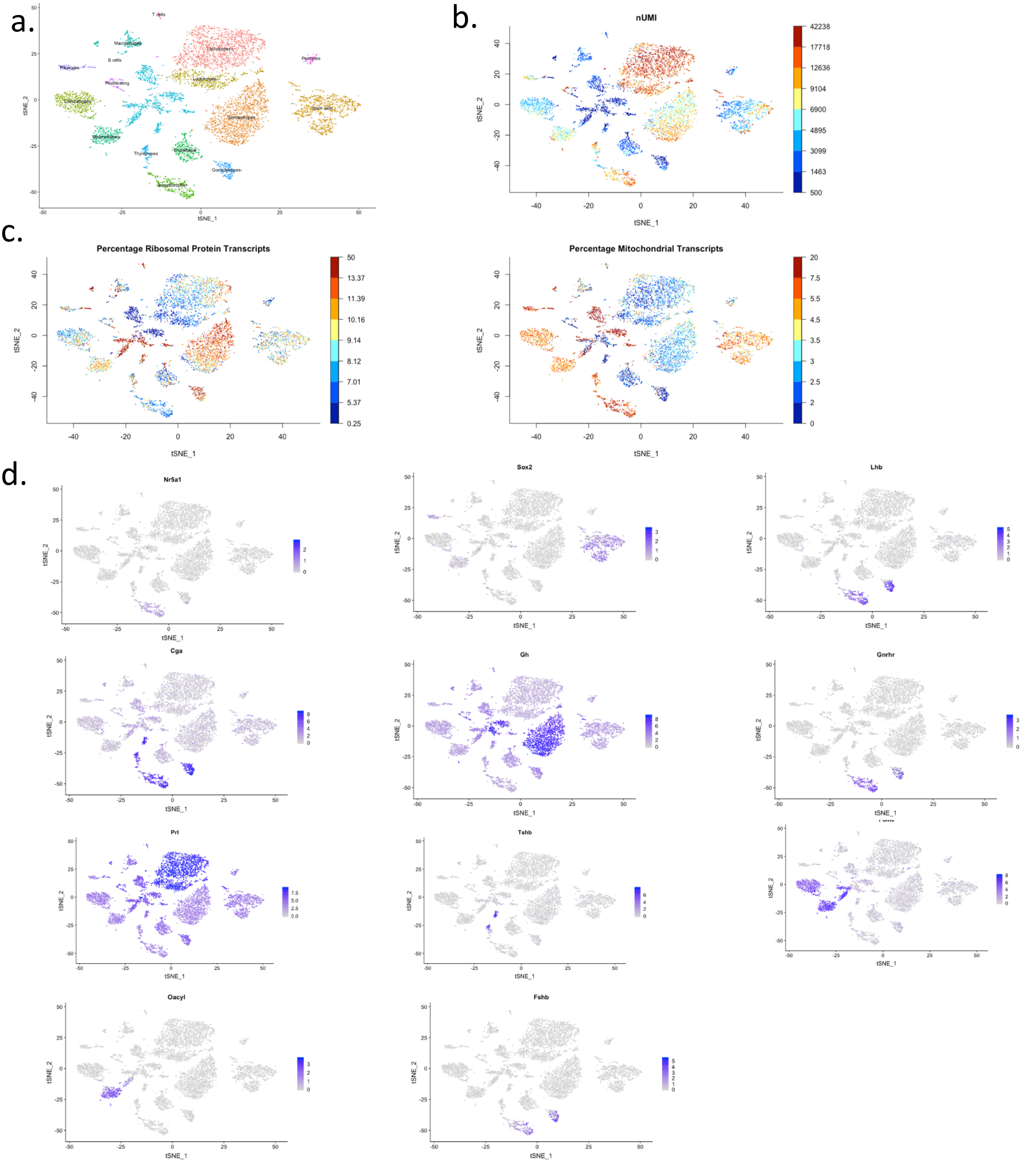


**Supplementary Figure 5:**
ScRNAseq analysis of a gently dissociated female pituitary. **a**. tSNE showing the different clusters obtained. **b**. Analysis of UMI content. **c**. Analysis of ribosomal content (Left panel), and mitochondrial content (Right panel). **d**. Feature plots showing the main pituitary markers for cell identification. A similar analysis was repeated in another dissociated female, whereas in dissociated male samples a different protocol was used (refer to **Supplementary Figure 1**). See **Supplementary Figure 6** for the same sample analyzed by snRNAseq, and **Supplementary Figure 7** for the same sample analyzed by snATACseq.

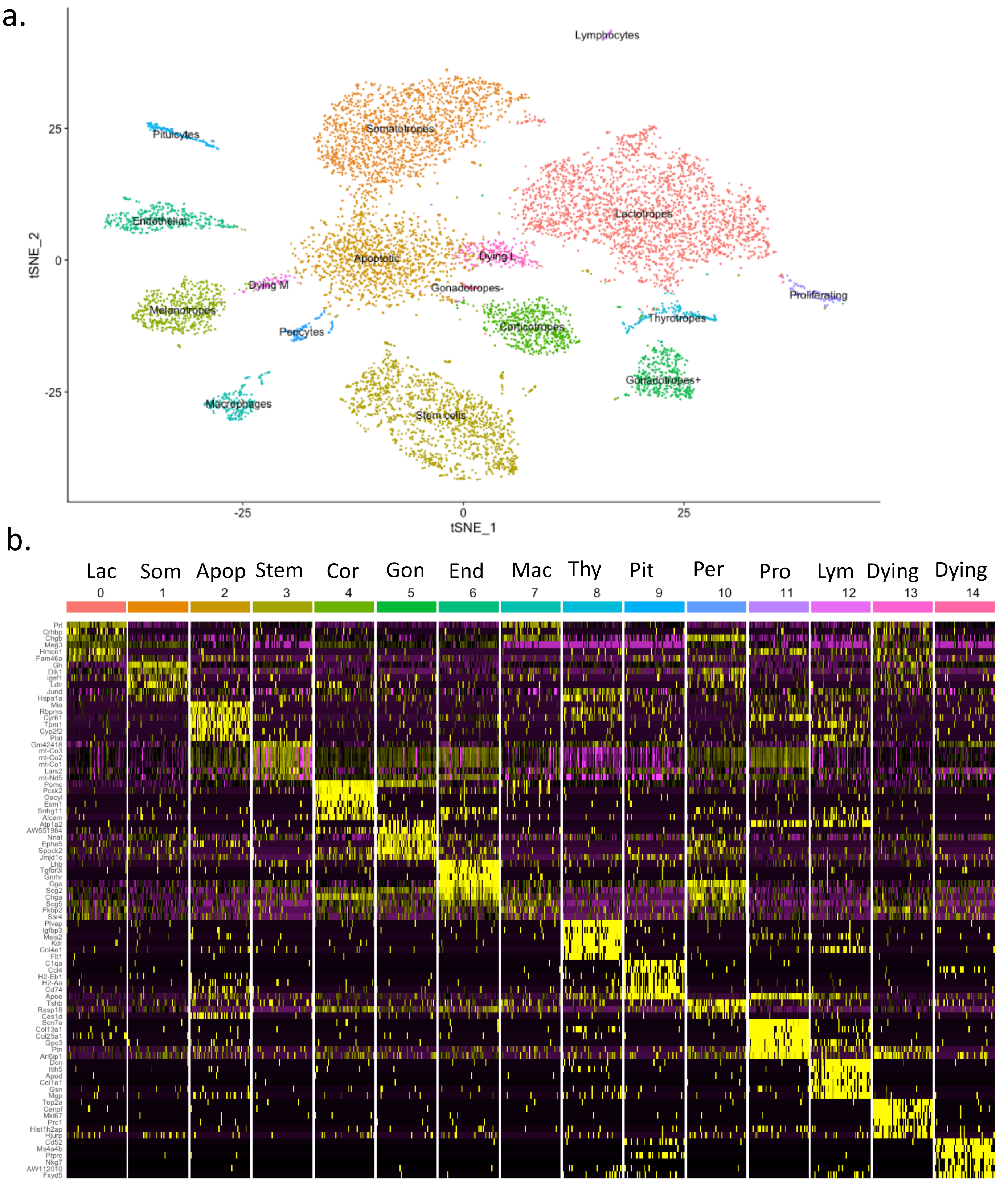


**Supplementary Figure 6:**
SCRNAseq analysis of a gently dissociated female pituitary (refer to **Supplementary Figure 5** for the same sample analyzed by scRNAseq). **a**. tSNE showing the different clusters that were obtained. **b**. Heatmap of showing cell identity based on gene expression in the sample. Note that contrary in contrast to scRNAseq (**Supplementary Figure 5**) we do not see the low UMIs cell clusters with snRNAseq.

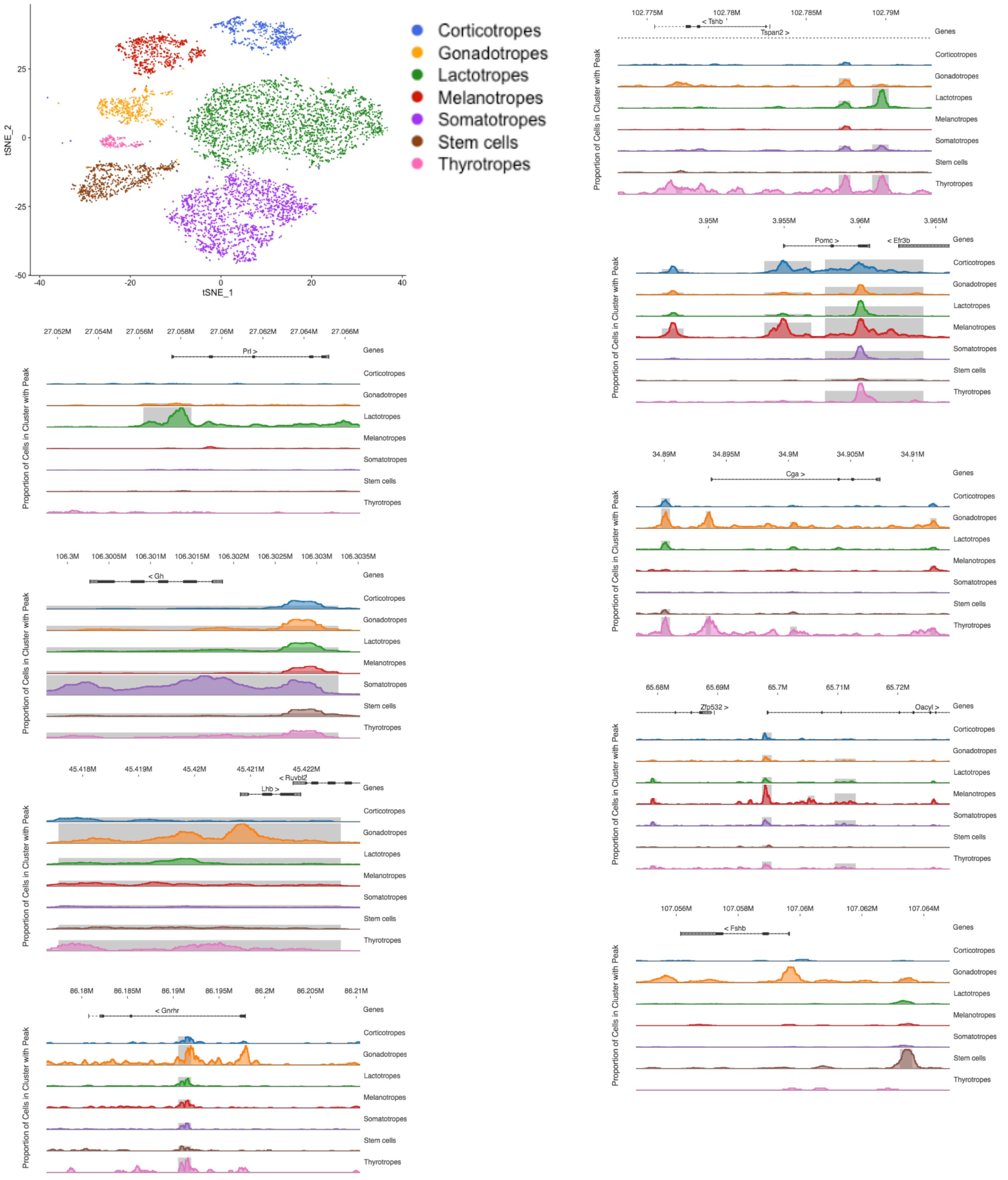


**Supplementary Figure 7:**

SnATACseq analysis of a gently dissociated female pituitary (refer to **Supplementary Figure 5** for the same sample analyzed by scRNAseq, and **Supplementary Figure 6** for the same sample analyzed by snRNAseq). **a**. tSNE identifying the different clusters that were obtained. **b**. Feature plots showing chromatin accessibility for several pituitary markers in the different cell-type clusters.

a.

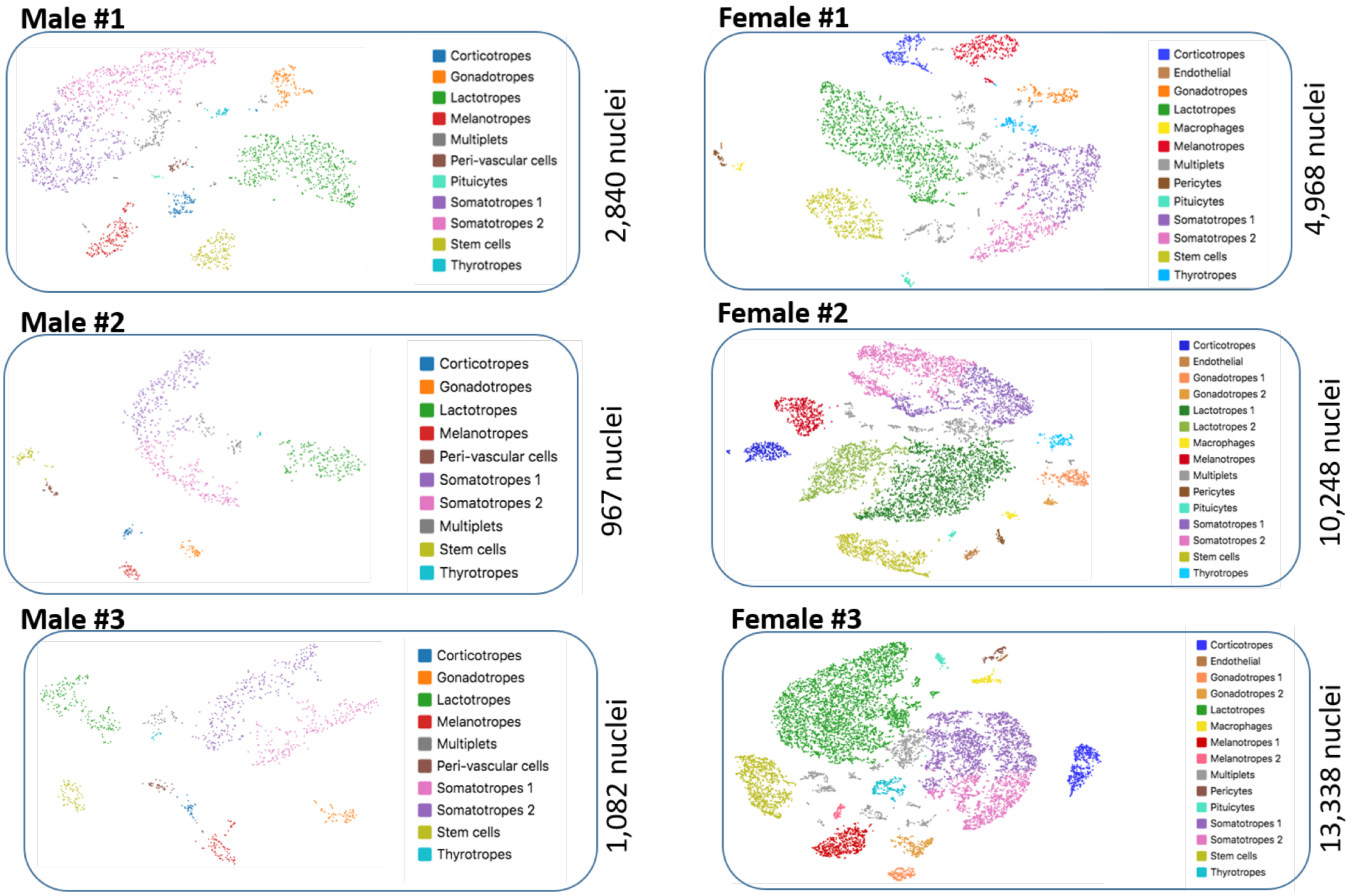


b.

| | FM1 | FM2 | FM3 | MM1 | MM2 | MM3 | Average |
|---|---|---|---|---|---|---|---|
| **Number of cells** | 4,968 | 10,248 | 13,338 | 2,840 | 967 | 1,082 | 5,574 |
| **Median fragments/cell** | 22,722 | 15,329 | 11,740 | 34,043 | 44,182 | 48,446 | 29,410 |
| **Fraction of fragments overlapping any targeted region** | 74.90% | 68.60% | 71.60% | 71.80% | 76.10% | 71% | 72% |
| **Fraction of transposition events in peaks in cell barcodes** | 73.20% | 65.70% | 65.80% | 65.40% | 75.60% | 67.80% | 68.92% |
| **Total number of read pairs** | 342,097,953 | 321,522,809 | 345,158,674 | 313,157,573 | 318,016,006 | 397,789,926 | 339,623,824 |
| **Fraction of reads with a valid barcode** | 96.40% | 96.10% | 96.10% | 96.10% | 96.50% | 96.40% | 96.27% |
| **Enrichment score of transcription start sites** | 11.36 | 8.26 | 9.58 | 9.92 | 11.82 | 9.08 | 10.0033333 |
| **Percent duplicates** | 44.50% | 32.70% | 32.20% | 45% | 63.40% | 59.70% | 46.25% |
| **Fraction of fragments overlapping called peaks** | 74.70% | 67.20% | 67.50% | 68% | 77.20% | 69.40% | 70.58% |

**Supplementary Figure 8:**
Results obtained for snap-frozen pituitaries that were processed for snATACseq. **a**. tSNE of all individual animals processed for snATACseq from individually snap-frozen pituitaries (3 males and 3 females). **b**. Metrics obtained on all 6 samples. These are the same samples that were processed for snRNAseq (**Supplementary Figure 2**).

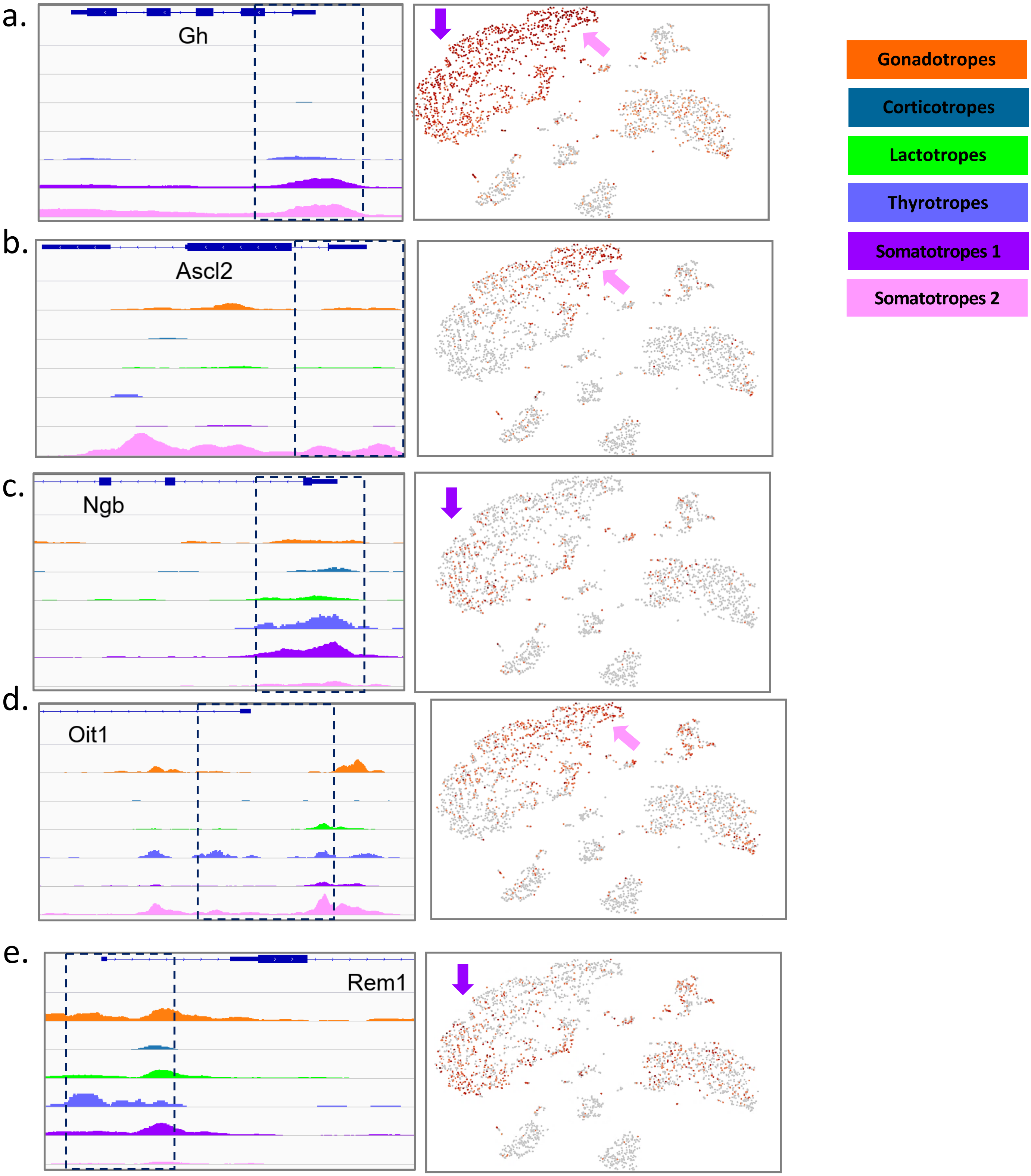


**Supplementary Figure 9:**
Analysis of the somatotrope gradient cluster by snATACseq illustrating **a**. homogeneous *Gh* promoter accessibility throughout the cluster; **b**. accessibility of the *Ascl2* promoter at the Som 2 pole; **c**. accessibility of the *Ngb* promoter at the Som 1 pole; **d**. accessibility of the *Oit1* promoter at the Som 2 pole; **e**. accessibility of the *Rem1* promoter at the Som 1 pole.

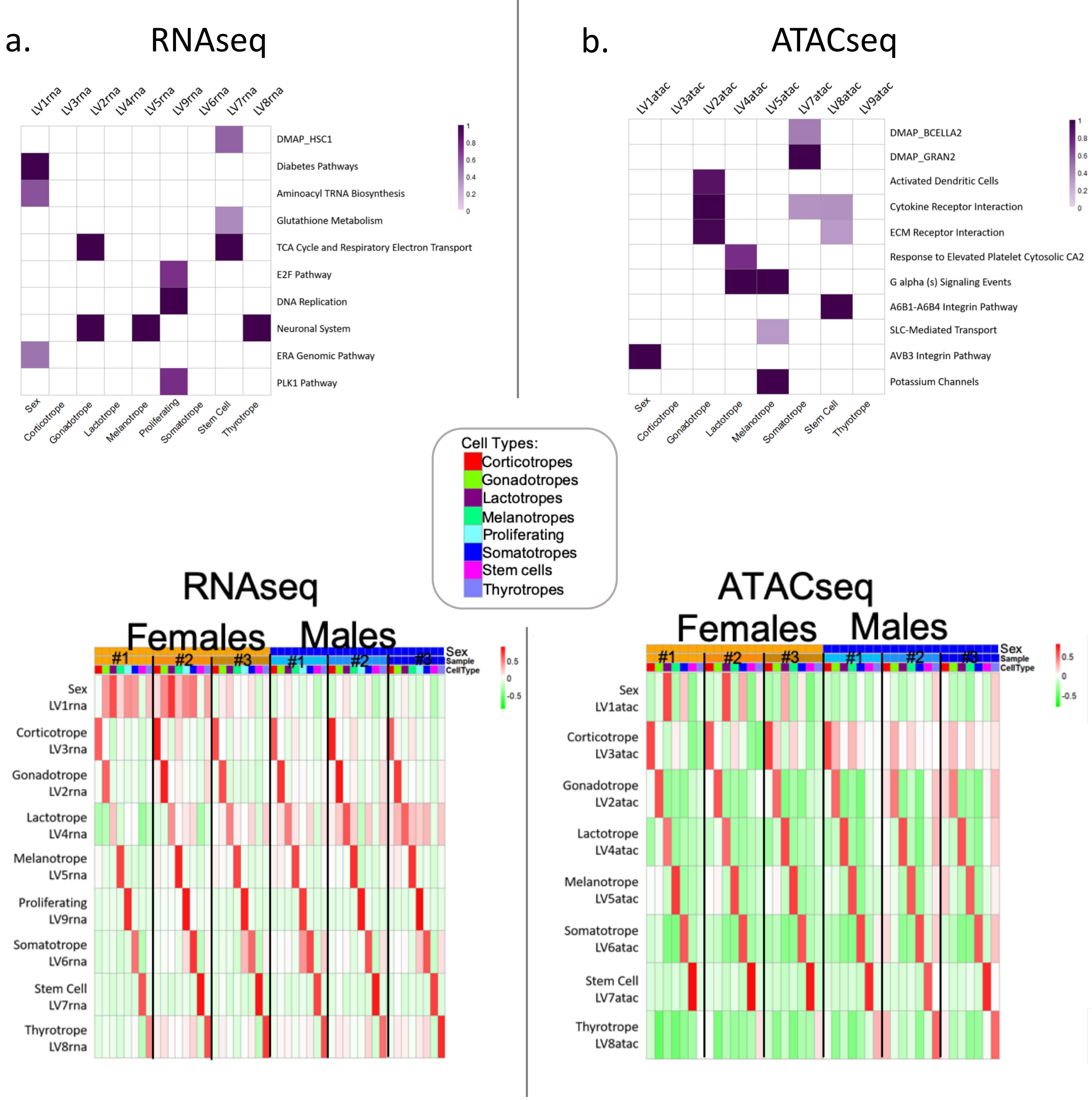


**Supplementary Figure 10:**

**a,b**. PLIER analysis of the snRNAseq (**a**.) and snATACseq data (**b**.) from the six individual snap-frozen pituitaries. Top panel shows the pathways associated with each sex or cell type-specific LV obtained from the given PLIER analysis. Heatmap value reflects the AUC of a cross-validation classification test of pathway association. Bottom panel shows a heatmap of the level of expression of each LV for a specific animal/cell type. Refer to **Fig. 6**.

### LV1rna, sex differences

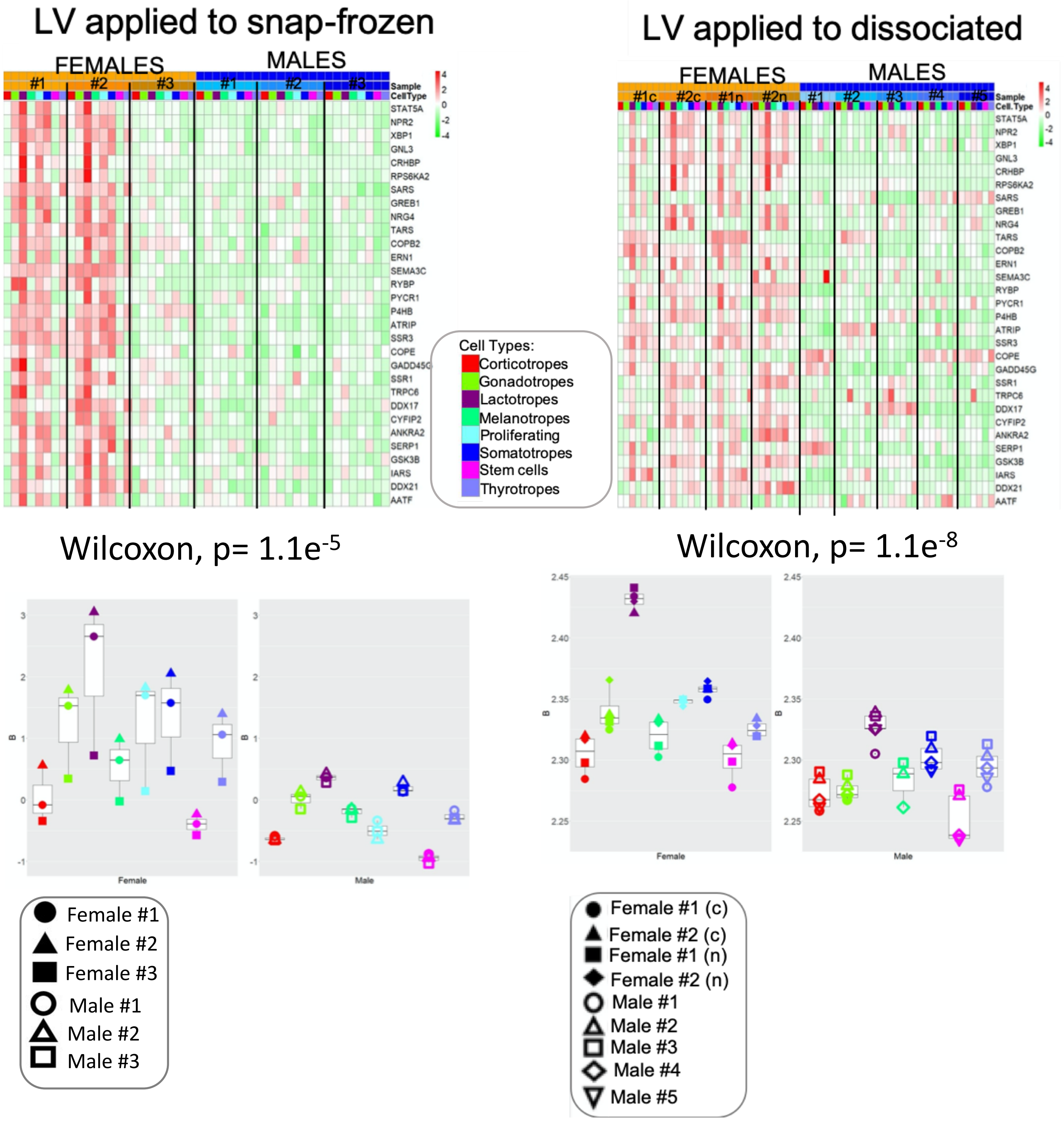


**Supplementary Figure 11:**
PLIER analysis showing comparison of the sex differentiating LV (LV1rna) in snap-frozen pituitaries (Left panels), vs. in dissociated and cryopreserved pituitaries (Right panels). Top panels: heatmaps showing the top 30 genes of the sex differentiating LV1rna. Bottom panels: boxplots showing the statistical analysis. LV1 RNAseq showed increased expression in females compared to males in the snap-frozen pituitaries (Wilcoxon $p=1.1e^{-5}$ ; refer to **Fig. 6a**), as well as in the dissociated pituitaries (Wilcoxon $p=1.1e^{-8}$). Dissociated female pituitary sample were analyzed by scRNAseq (c) and snRNAseq (n). Dissociated male pituitary samples were analyzed in scRNAseq only.

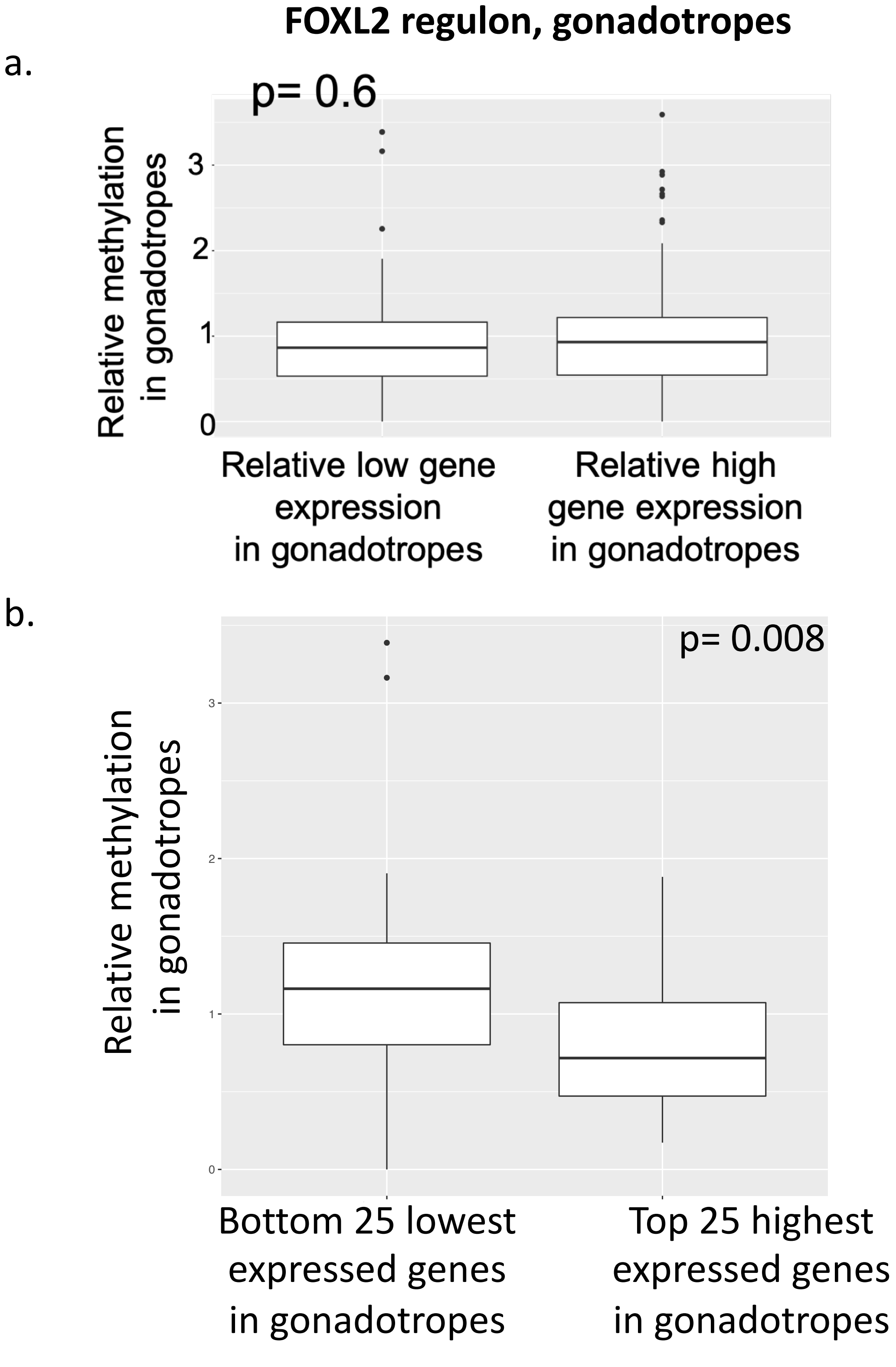


**Supplementary Figure 12:**
**a**. Boxplots of relative methylation at the TSS for all 331 genes composing the FOXL2 regulon, separated into relatively low expression vs. relatively high expression in male gonadotropes (consensus male from 3 individual male samples). **b**. Boxplots of the relative methylation at the TSS for the 25 lowest and 25 highest expressed genes of the FOXL2 regulon in male gonadotropes (consensus male from 3 individual male samples). Refer to **Fig. 7**.

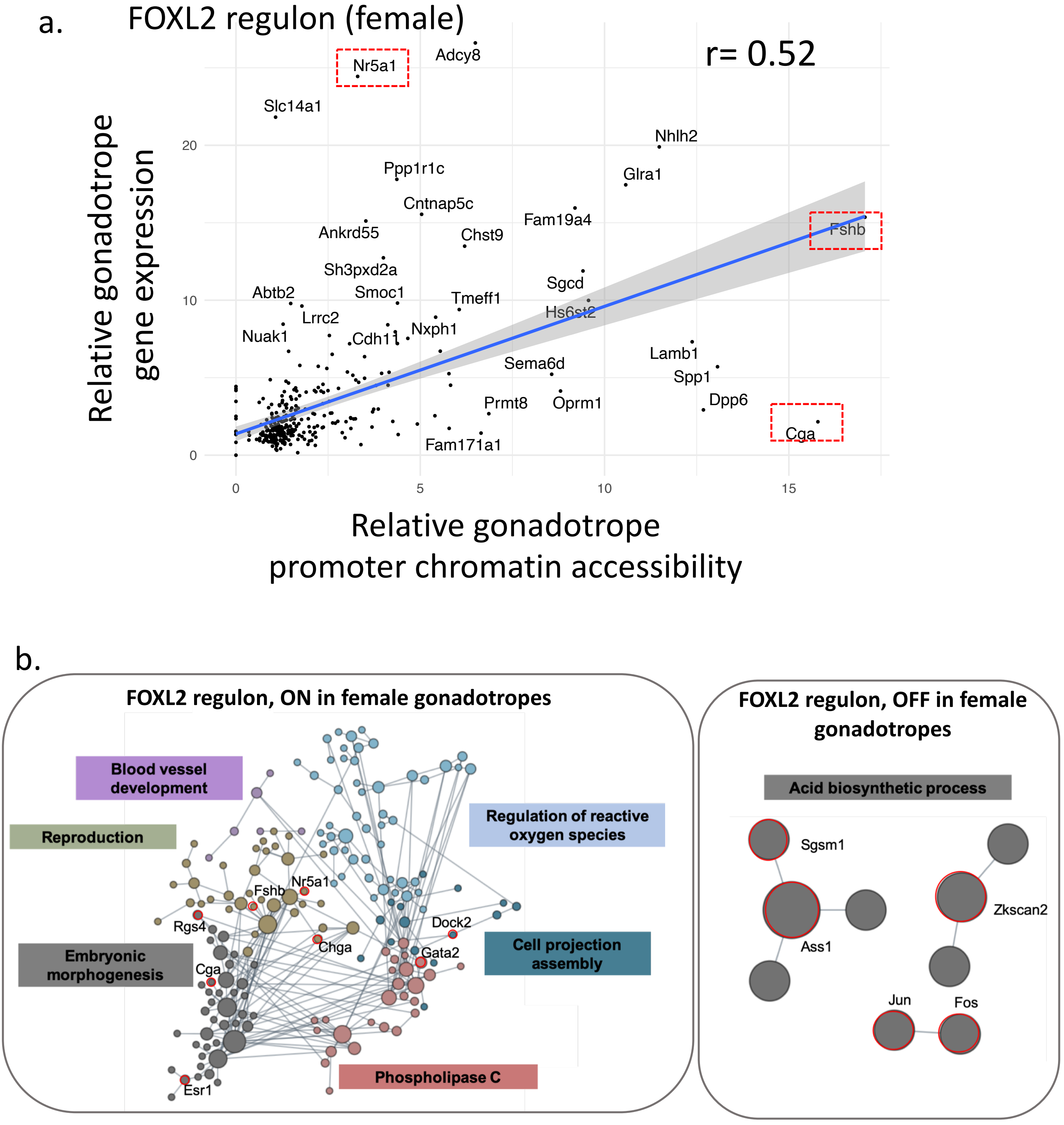


**Supplementary Figure 13:**
**a**. Scatter plot and correlation of the 331 genes composing the FOXL2 regulon based on their relative gene expression and relative TSS chromatin accessibility in female gonadotropes (consensus female from 3 individual female samples). Boxed in red are genes that are related to gonadotrope regulation. Refer to **Fig.7D** for males. **b**. Gene network for the FOXL2 regulon targets that are ON in female gonadotropes (Left panel), and regulon targets that are OFF in female gonadotropes (Right panel; consensus female from 3 individual male samples).

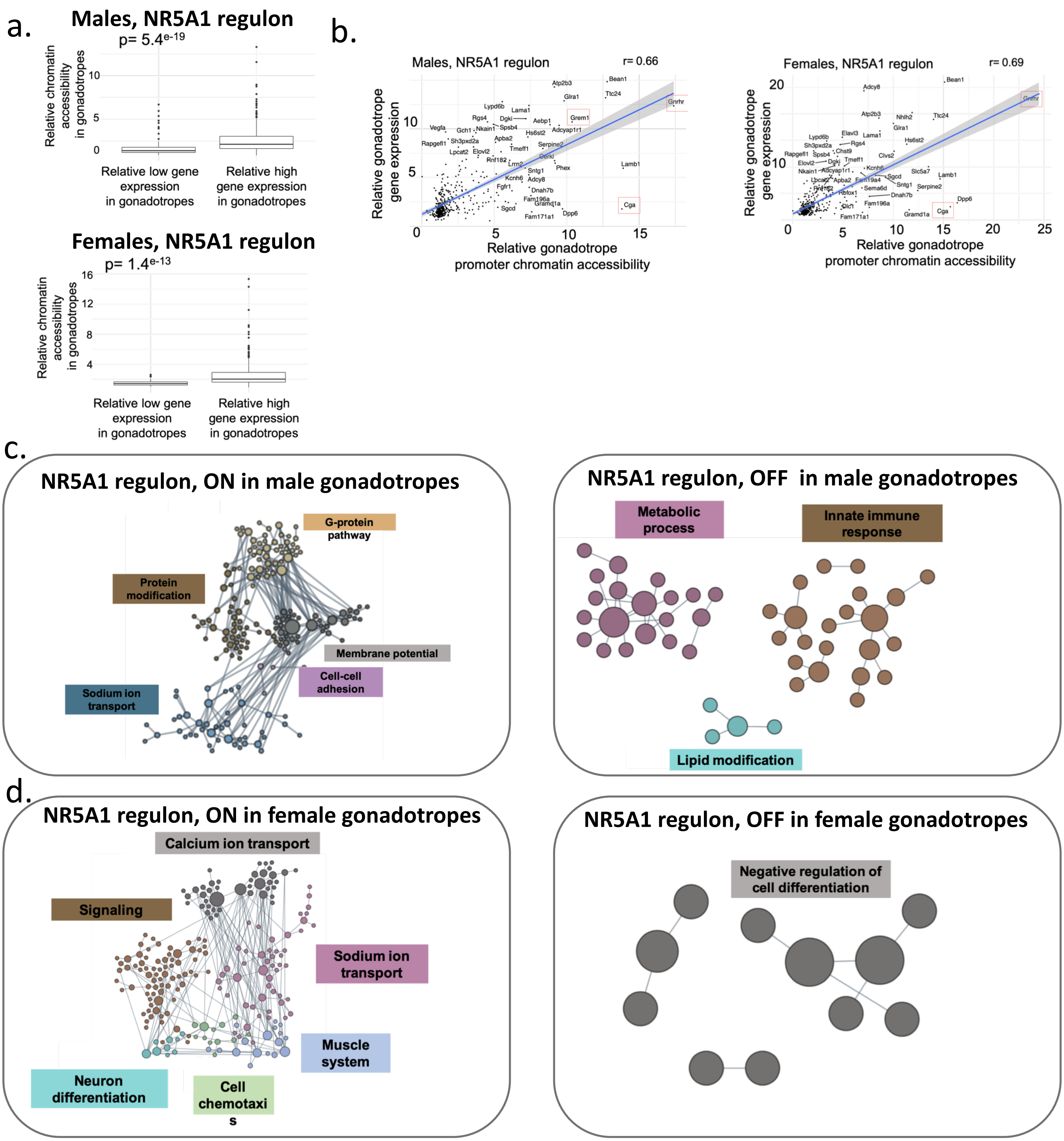


**Supplementary Figure 14:**

**a**. Boxplots of the relative chromatin accessibility within gene bodies for all 412 genes composing the NR5A1 regulon separated into relatively low expression vs. relatively high expression in male (Top panel) and female (Bottom panel) gonadotropes. **b**. Scatter plot and correlation of the 412 genes composing the NR5A1 regulon based on their relative gene expression and relative TSS chromatin accessibility in male (Left panel) and female (Right panel) gonadotropes. Boxed in red are genes that are related to gonadotrope regulation. **c**. Gene network for the NR5A1 regulon targets that are ON (Left panel), and regulon targets that are OFF in male gonadotropes (Right panel). **d**. Gene network for the NR5A1 regulon targets that are ON (Left panel), and regulon targets that are OFF in female gonadotropes (Right panel). All panels are consensus from 3 animals.

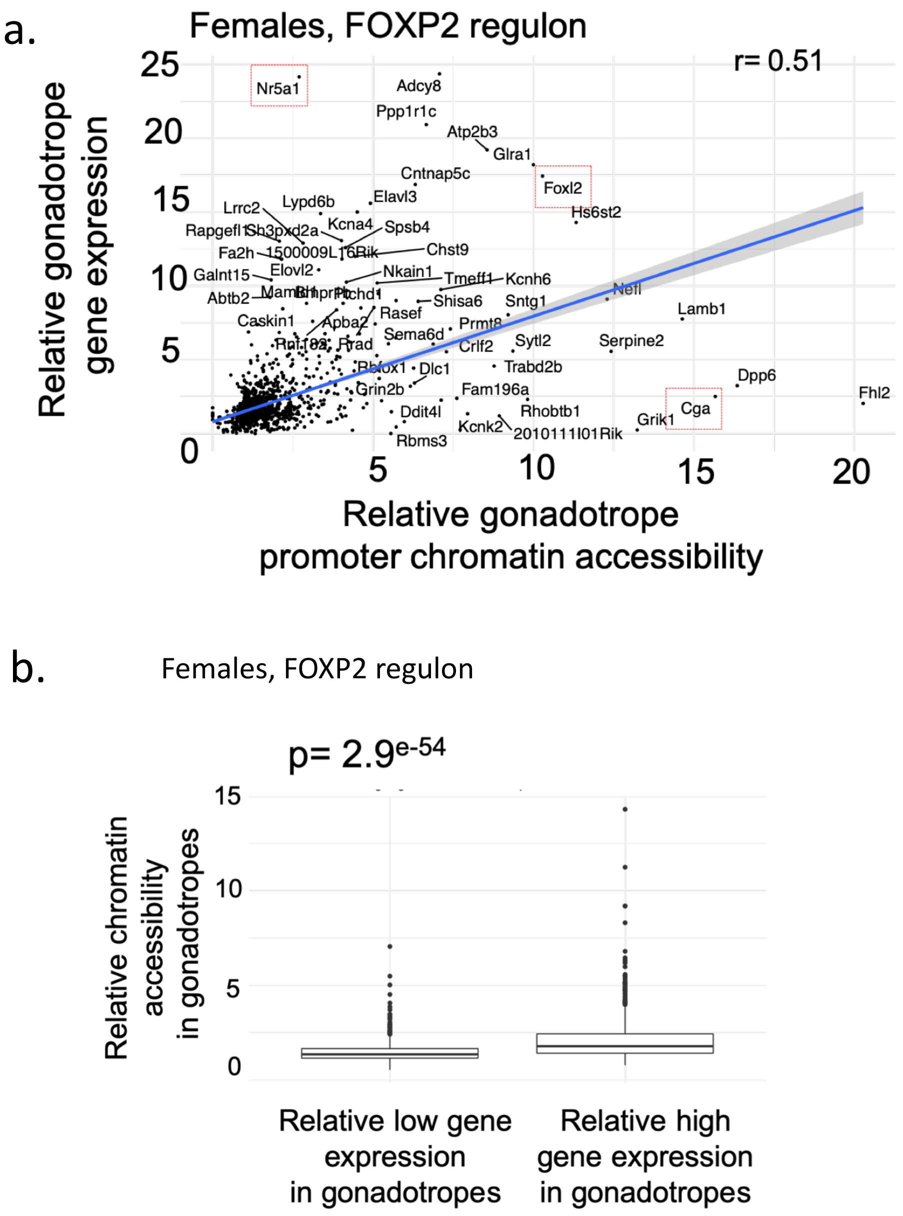


**Supplementary Figure 15:**
**a**. Scatter plot and correlation of the 1676 genes composing the FOXP2 regulon based on their relative gene expression and relative TSS chromatin accessibility in female gonadotropes (consensus from 3 individual female samples). Boxed in red are genes that are related to gonadotrope regulation. **b**. Boxplots of the relative chromatin accessibility within gene bodies for the 1676 genes composing the FOXP2 regulon separated into relatively low expression vs. relatively high expression in female (right) gonadotropes (consensus from 3 individual female samples). Refer to **Fig. 8** for analysis of the FOXP2 regulon in males.

### *Fshb*

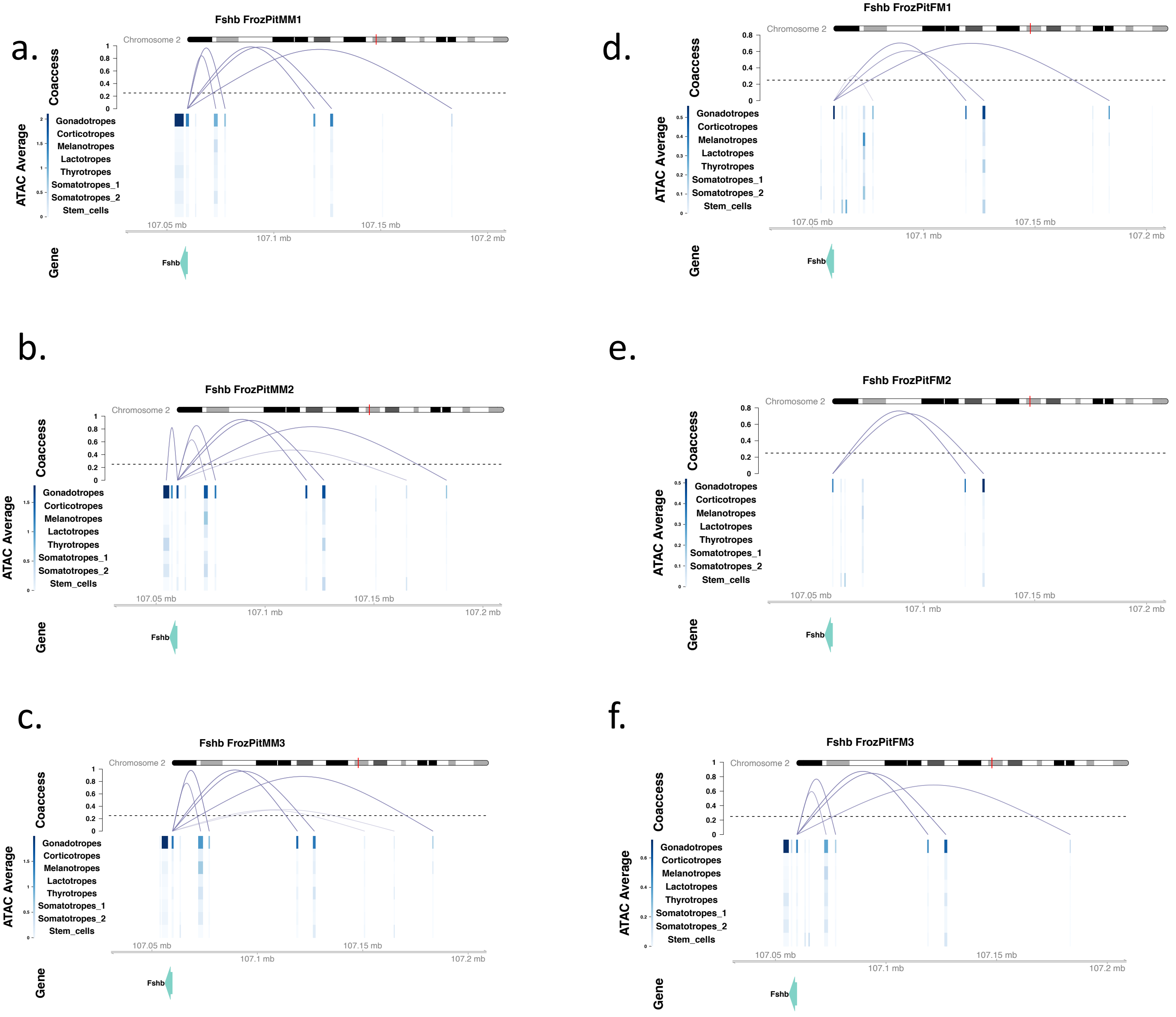


**Supplementary Figure 16:**
Co-accessibility analysis for the *Fshb* promoter in all 6 individual snap-frozen pituitaries (3 males (**a-c**) and 3 females (**d-f**)). Refer to **Fig. 9a**.
